# Alzheimer disease—The proposed role of tanycytes in the formation of tau tangles and amyloid beta plaques in human brain

**DOI:** 10.1101/2025.04.08.647836

**Authors:** Ruth Fabian-Fine, Abigail G. Roman, Adam L. Weaver

## Abstract

The causes of Alzheimer Disease (AD) are attributed to the formation of abnormal protein deposits including tau tangles and Amyloid ß (Aβ) plaques. These proteins are believed to accumulate in the cortex due to impaired waste removal resulting in neurodegeneration. Interestingly, removal of these protein accumulations has failed to halt disease progression. How can this be explained? We have recently proposed the existence of a tanycyte-derived canal-system in the brain that likely internalizes neuronal waste. We propose that Aβ and tau protein may play important structural roles in this canal system that swells in AD-affected brain. Using *in situ* hybridization, immunohistochemistry, ultrastructural, and histological methodologies, here we demonstrate the formation of receptacles from translucent ‘swell-bodies’ that form along tanycyte processes. We show that both spatial distribution and anatomical appearance of swell-bodies differ from myelin-forming ependymal tanycytes. Interestingly, receptacles that differentiate within swell-bodies express mRNA for aquaporin4 and the AD-related proteins Presenilin1 and amyloid precursor protein. Consistent with these findings we show strong Aβ immunolabeling of these receptacles that swell and proliferate in AD-affected brain and take on the appearance of Aβ plaques. We conclude that it is likely pathological abnormalities of the associated tanycytes leads to this abnormal swelling.

## Introduction

Currently, the underlying causes for Alzheimer Disease (AD) are elusive and are thought to lie in the formation of abnormal intracellular deposits of hyperphosphorylated tau tangles and extracellular Amyloid ß (Aß) plaques throughout the human cortex. Due to this abnormal protein accumulation in AD-affected brain tissue, a prevalent causative hypothesis is failed waste removal from the nervous system [1–4]. Currently, the way by which waste is cleared from the brain is poorly understood. Illiff et al. (2012) have proposed the existence of an aquaporin-4 aqua channel (AQP4)-mediated ‘Glymphatic System.’ Based on this postulation AQP4-expressing astrocytes are thought to flush cellular debris from the brain by means of a cytoplasmic convective flow of cerebrospinal fluid. Using two-photon microscopy and fluorescent tracers in mouse brain, the authors demonstrated the formation of a convective flow through the neuropil believed to flush cellular waste into peri-venous spaces for clearance from the brain [1, 5–7]. Interestingly, a significantly reduced flow of cerebrospinal fluid was observed in AQP4-null mice compared to wild-type mice [8]. However, this hypothesis has been put into question predominantly regarding the proposed astrocyte-induced AQP4-mediated flow of brain fluids [3, 9]. In an alternative postulation, recent studies have suggested that hypothalamic and hippocampal ependymal tanycytes may mediate waste removal from the brain [10, 11]. Interestingly in both studies, structural tanycyte pathologies were observed in AD affected brain tissue. Sauvé et al. [12] have demonstrated the comparatively rapid clearance of tau protein from cerebrospinal fluid into the blood stream within 30 min. The authors propose that tanycyte fragmentation in AD impairs this mechanism, leading to accumulation of tau protein in affected patients. We have previously demonstrated that myelinated ependymal tanycyte processes in the human hippocampus form varicosities consistent with observations in hypothalamic tanycytes as reported by Pasquettaz et al., (2021) [13]. We have shown, at both light- and electron microscopic levels, that hippocampal tanycytes form myelin-derived protrusions that project into neuronal somata and differentiate into waste-internalizing receptacles consistent with the appearance of waste-containing lipofuscin [10, 14]. Currently, the main functional roles of tanycytes are the proposed metabolic and energetic support of neurons in addition to the formation of tight junctions [15–21]. However, considering these new findings we suggest that tanycytes may also play an important role in waste clearance from the brain.

Guided by the recently discovered, myelin-derived waste canal system in the Central American wandering spider *Cupiennius salei* [10, 22], we have proposed that debris uptake in the hippocampal formation may be mediated by a similar ‘glial-canal system’ [10]. In giant spider neurons, these myelin-derived waste-containing canals are clearly visible due to their large sizes. Because of this advantage, we discovered that this canal system is likely formed through the synergistic interaction of two cell types. Firstly, large myelin-forming macroglia that reside around the periphery of individual ganglia (equivalent of functional nuclei in the mammalian brain) and form a vast interconnected myelin-network. This network sends long myelinated processes into the brain parenchyma and form projections into each neuron [10, 22]. The second cell type forms long slender processes with translucent lumina that transect into the myelinated cells and neurons. We have demonstrated that AQP4-immunoreactive cell profiles that form large translucent processes are often located within the lumina of myelin-derived waste canals (see also summary Fig 12 here). This location of the AQP4-expressing cell profiles within the myelin-derived canal structures would be highly suitable to create a convective bulk flow toward the myelin-derived canals. This would explain why vast amounts of lipofuscin accumulate around and within these glial canals ([22]; see also summary figure below). As demonstrated here, we postulate that this synergistic, myelin-AQP4 derived canal system is highly conserved in rodent and human brain and internalizes waste for removal from the hippocampal parenchyma. We propose a similar synergistic interaction of myelin-forming and AQP4-expressing ependymal macroglia, which we propose to be tanycytes. Consistent with findings by Sauvé et al. [11], we have provided evidence that the canal-forming tanycytes show structural pathologies in AD-affected brain tissue. In hippocampal tanycytes, these pathologies are consistent with hypertrophic swelling [10]. Here, we have investigated mouse, rat, and human hippocampus and provide additional histological, ultrastructural, and biochemical evidence that is consistent with the proposed existence of a waste-internalizing glial-canal system. We demonstrate that specialized waste-containing receptacles emanate from this canal system and project into both extracellular spaces and neuronal somata. We show that these receptacles express the Aß-related genes for presenilin-1 (Pres1) and amyloid precursor protein (APP), which is consistent with their Aß-immunolabeled nature in non-AD affected human hippocampus. We furthermore demonstrate that these receptacles swell in AD-affected brain tissue and resemble Aß-plaques [23] in immunolabeled brain sections. Our findings are consistent with the previously proposed failure of waste removal in AD affected brain [24, 25]. However, based on the findings presented and discussed here we propose an alternative, tanycyte-mediated mechanism by which waste is removed from the brain. As discussed below, our hypothesis provides a compelling explanation for the prevalence of Aß and tau aggregates in AD-affected human brain and why removing these proteins from AD-affected brain does not halt disease progression.

## Materials & Methods

| Key Resources Table |  |  |  |  |
| --- | --- | --- | --- | --- |
| Reagent type (species) or resource | Designation | Source or reference | Identifiers | Additional information |
| *Mus musculus* | C57Bl6 | University of Vermont Animal Facilities | Heppner et al 2024 | Surplus brain tissue |
| *Rattus norvegicus*: Sprague-Dawley adult brain blocks | NCBI:txid10116 | Univ. College London |  | doi: 10.1016/s0896-6273(03)00559-2 |
| antibody | Cy3-goat-anti rabbit | Jackson ImmunoResearch | Cat# 111-165-003, RRID:AB_2338000 | 1:1000 Mouse uptake experiments |
| antibody | Alexa 488-donkey anti-goat | Jackson ImmunoResearch | Cat# 705-545-003, RRID:AB_2340428 | 1:1000 Mouse uptake experiments |
| AQP4 channel blocker | TGN-020 | Sigma Aldrich | SML0136 | 60 µm concentration |
| Fixative | Paraformaldehyde | Electron Microscopic Sciences | EMS 15710 | 4% Freshly prepared |
| Fixative | Glutaraldehyde | Electron Microscopic Sciences | EMS 16019 | 2.5% freshly prepared |
| Fixative | Osmium tetroxide | Electron Microscopic Sciences | EMS 19150 | 1% Solution |
| Tissue embedding | Agarose | Sigma Aldrich | Cat# A 9539 | 4% Concentration |
| antibody | rabbit anti-aquaporin-4 | BiCell | Cat# 20104 | Immunohistochemistry (1:100) |
| antibody | anti-Caspase2 | BiCell | Cat# 10302 | Immunohistochemistry (1:100) |
| antibody | anti-Pres1<br>(rabbit monoclonal) | Sigma Aldrich | Cat#: ZRB1614 | Immunohistochemistry (1:100) |
| antibody | anti-APP<br>(mouse monoclonal) | Sigma Aldrich | Cat#: MABN380,<br>RRID:AB_2714163 | Immunohistochemistry (1:100) |
| antibody | anti-alpha tubulin (mouse monoclonal) | DSHB IOWA | Cat#: mouse anti-α-tubulin 12G10,<br>RRID:AB_1157911 | Immunohistochemistry (1:100) |
| antibody | anti-myelin<br>(mouse monoclonal) | DSHB IOWA | Cat#: mouse anti-Myelin 6-4H2,<br>RRID:AB_2877129 | Immunohistochemistry (1:100) |
| antibody | Cy3 goat anti-rabbit | Jackson ImmunoResearch | Cat# 111-165-003 | Immunohistochemistry 1:600 |
| antibody | Cy3 goat anti-mouse | Jackson ImmunoResearch | Cat# 115-165-148 | Immunohistochemistry 1:600 |
| antibody | Cy3 goat anti-rat | Jackson ImmunoResearch | Cat# 112-165-003 | Immunohistochemistry 1:600 |
| Nuclear stain | Hoechst Blue | Sigma Aldrich | 94403-1ML | 1:3000 |
| antibody | mouse anti-AT8 | Thermo Fisher Scientific | Cat# MN1020 | Immunohistochemistry 1:1600 |
| antibody | mouse anti-AT8 | Thermo Fisher Scientific | Cat# MN1020 | Immunohistochemistry 1:1600 |
| antibody | mouse anti-AT8 | Abcam | Cat# ab20106 | Immunohistochemistry 1:1000 |
| Histological myelin stain | Luxol Fast Blue | Alfa Aesar | A15395 | 0.1% concentration |
| mRNA complementary probe | Aquaporin 4 | Bio-Techne | #482448 | 1x dilution |
| mRNA complementary probe | Myelin Basic Protein | Bio-Techne | #411058-C2 | 1x dilution |
| mRNA complementary probe | Pan-Quaking | Bio-Techne | #416358-C3 | 1x dilution |
| mRNA complementary probe | Glial Fibrillary Acid Protein | Bio-Techne | #311808-C2 | 1x dilution |
| mRNA complementary probe | Sox10 | Bio-Techne | #484121-C4 | 1x dilution |
| mRNA complementary probe | Caspase3 | Bio-Techne | 436568-C3 | 1x dilution |

### Mouse brain tissue

The mouse brain originated from four adult (6-12 month-old) male (2) and female (2) house mice (*Mus musculus*) of the GCaMP6f UPII-Cre strain. The animals were raised at the University of Vermont animal facilities. These mice were wildtype mice regarding neuronal and glial cells in the brain and were raised for bladder-related studies [26]. For ethical reasons and to avoid unnecessary animal sacrifices, we have utilized the brains of these animals for our preliminary ‘test of concept’ experiments. The tissue was obtained freshly upon dissection of the animals and processed as described below. IACUC approval was granted under PROTO202200018 (University of Vermont) and IACUC-2024-001 Fabian-Fine (Saint Michael’s College).

### Rat brain tissue

For ethical reasons to avoid unnecessary animal sacrifices, the hippocampus used in this study originated from healthy adult male Sprague Dawley control rats previously processed for electron-microscopy during prior studies on rat hippocampus carried out by the first author. All experiments were conducted in accordance with prevailing regulatory animal protocols in 2003 at University College London. The tissue embedding for semi- and ultra-thin sections was carried out as described below

### Human brain tissue

The human brain tissue used for our investigations originated from five AD-affected hippocampal autopsy brain samples (86 y/o male A2B2C1 intermediate Alzheimer’s disease neuropathologic change (ADNC); 86 y/o female A3B3C2 high ADNC;79 y/o male A3B2C1 intermediate ADNC; 96 y/o male A3B2C2 intermediate ADNC; 86 y/o female A3B3C3 high ADNC) and five hippocampal autopsy brain samples that did not show signs of ADNC (33 y/o male; 40 y/o male; 53 y/o female; 39 y/o male; 38 y/o female). The tissue was fully consented, including ‘consent for research,’ from next of kin in compliance with Vermont State law, the Health and Human services regulation 45 CFR 46.102(e), and the University of Vermont regulations regarding human subject research. The autopsy examination was carried out by Dr. John DeWitt at the University of Vermont Medical Center.

### Uptake experiments on living mouse brain

Three mice from which the surplus brain tissue was obtained were euthanized with Euthasol (Virbac) by a licensed technician. The brains were immediately removed from the cranium and transferred into mouse culture medium DMEM (VWR Cat #45000-312) containing 4.5 g/L glucose, L-glutamine, 10% Fetal Bovine Serum and 1% Penicillin/Streptomycin at 37 °C to maintain temperature-appropriate enzymatic activity. The two brain hemispheres were separated, and each hemisphere was sliced into three equal coronal sections along the anterior-posterior axis. The sections from the right hemisphere (controls) were kept in culture medium, whereas the left hemisphere sections (experimental) were kept in culture medium to which we added Cy3-coupled goat-anti rabbit secondary antibody (1:1000; Jackson ImmunoResearch # 111-165-003; RRID:AB_2338000). After 30 min, both control and experimental brain sections were fixed with freshly made 4% paraformaldehyde (4% PFA) overnight at 6 °C. The preparations were embedded in 4% agarose (Sigma 9539) and sectioned into 70-µm vibratome sections using a Leica Vibratome 1000S. The sections were washed in PBS and mounted on glass slides using Mowiol (Sigma 81381). The sections of both control and experimental preparations were imaged with identical settings using a Zeiss Axio-Imager confocal microscope with ZEN-Blue software. To test that the internalized fluorochrome detected in ependymal cells of the experimental preparations indeed originate from the Cy3-coupled goat antibody, we utilized a secondary Alexa 488-coupled donkey anti-goat antibody (Jackson ImmunoResearch # 705-545-003; RRID:AB_2340428). For this purpose, additional sections were cut and washed in 0.1 M phosphate-buffered saline pH 7.4 (PBS). Unspecific binding sites were blocked with blocking medium containing 0.25% Bovine Serum Albumin (Sigma A4503) and 5% Normal goat serum (Sigma G9023) in 1% Triton-X/PBS for 20 min prior to incubation with the donkey anti-goat antibody (1:600 overnight at 6 °C). Subsequently, the preparations were washed in PBS (2×5 min), stained with Hoechst Blue nuclear stain (1:3000; 15 min), washed in PBS 5×10 min, and mounted on glass slides using Mowiol prior to examination with the Zeiss confocal microscope.

### Image and statistical analysis

Confocal images were exported from the Zeiss ZEN-Blue program using the “Original Data” setting and analyzed in CellProfiler Image Analysis Software 4.2.8 (RRID:SCR_007358) for all measurements [27]. Briefly, Cy3 stained objects were identified [Three-class Otsu thresholding; middle class: background] that were brighter than the background and had a diameter between 5 and 60 µm. Object size/shape and intensity were measured. In some cases, full captured images had their exposure minimally modified to enhance contrast or brightness identically for figures, but not for the analysis.

An unpaired, two-tailed Student’s t-test with Welch’s correction was used to perform comparisons of the Log_10_ mean object intensities across control (0 µM) and TGN-020 AQP4-blocked (60 µM; m) conditions. All statistical parameters and results are reported in the relevant figure legend. To the best of our knowledge, all assumptions of this test were met. Statistics were calculated and graphs were created using GraphPad Prism 10.4.1 (RRID:SCR_002798).

### Tissue processing for semi- and ultrathin sectioning

Hippocampal and spider brain tissue samples (∼50 mm thick) were immersed in freshly prepared ice cold 4% paraformaldehyde (EMS 15710) and 2.5% glutaraldehyde (EMS 16019) in phosphate buffered saline (pH 7.4, 0.1 M; PBS) and fixed for at least 48 hours in the refrigerator. The preparations were embedded in 4% agarose and sectioned into 100 µm sections in PBS using a Leica 1000S Vibratome. The sections were postfixed in 1% osmium tetroxide (EMS 19150), dehydrated in a graded series of ethanol, and embedded into Araldite as described previously [10]. Using a Diatome Histo-knife with 8 mm blade ultrathin sections (50-60 nm) were cut with a Leica Ultracut E. Semithin sections were cut at 1 µm thickness, collected on glass slides, dried at 55°C on a hot plate, stained with filtered 1% aqueous toluidine blue solution for 1-3 min. After removal of excess stain with distilled water, preparations were dried and mounted in immersion oil to prevent destaining of the preparations.

Ultrathin sections were cut at 50-60 nm thickness. The sections were stretched with chloroform (Electron Microscopic Sciences 12540) by gently waving a chloroform-soaked wooden toothpick over the sections without touching them until they were sufficiently stretched. The sections were collected on pioloform-coated single-slot copper grids (EMS G2010CU) using #5 Dumont forceps and an eyelash that was mounted to a wooden toothpick using hot bee’s wax. Grids were contrasted with aqueous 1.5% uranyl acetate (6 min) and Reynold’s lead citrate (6 min) according to standard electron microscopic methods. The examination was conducted using a JOEL 1400 electron microscope with digital image acquisition operated at 80 kV.

### Light microscopic immunohistochemistry

Human hippocampal tissue utilized for light-microscopic immunolabeling was fixed in freshly prepared, ice-cold 4% paraformaldehyde for a minimum of 48 h at 6 °C. The tissue was embedded in 4% Agarose (Sigma # A 9539), sectioned into 70-µm vibratome sections using a Leica 1000S Vibratome. The sections were washed in PBS 4×5 min and unspecific binding sites were blocked with blocking medium containing 0.25% Bovine Serum Albumin (Sigma A4503) and 5% Normal goat serum (Sigma G9023) in 1% Triton-X/PBS for 20 min. The sections were incubated overnight at 6 °C within the respective primary antibody solutions (rabbit anti-aquaporin-4 antiserum BiCell 20104; rat anti-Caspase2 BiCell 10302; rabbit anti-Pres1, Sigma Aldrich ZRB1614; mouse anti-APP-C99, Sigma Aldrich MABN380, RRID:AB_2714163; mouse anti-α-tubulin 12G10, DSHB, IOWA, RRID:AB_1157911; and mouse anti-Myelin 6-4H2 DSHB, IOWA, RRID:AB_2877129) at dilutions of 1:100. Prior to incubation with the secondary antibodies, preparations were washed thoroughly in PBS (5×10 min), incubated in blocking medium as described above, and incubated with either secondary Cy3 goat anti-rabbit (Jackson ImmunoResearch Laboratories 111-165-003), Cy3 goat anti-mouse (Jackson ImmunoResearch # 115-165-148), and Cy3 goat-anti rat (Jackson ImmunoResearch # 112-165-003) at dilutions of 1:600 in PBS containing 10% blocking medium overnight at 6 °C. Subsequently, the sections were rinsed in PBS (3×2 min). For the nuclear stain, we utilized Hoechst Blue (Sigma Aldrich 94403-1ML) at a dilution of 1:3000 in PBS for 20 min. After washing in PBS (5×10 min), the sections were mounted on glass slides using Mowiol (Sigma 81381). To avoid bleaching of the fluorochromes, the mounting was conducted with minimum illumination.

### Toluidine-blue stain of vibratome sections for light microscopic investigations

Vibratome sections from 4% paraformaldehyde-fixed tissue were washed in PBS for four wash cycles. The toluidine blue staining procedure was conducted under visual control using a dissection stereo microscope by slowly dripping a 2% toluidine blue solution into a petri dish containing brain sections in PBS. The staining was terminated after approximately 2 min when the neurons appeared appropriately stained for light microscopic investigation without overstaining the preparations as this will result in the inability to distinguish individual structures. We utilized Mowiol as mounting medium. The preparations were examined promptly using an Olympus microscope with differential phase contrast and digital image capturing capabilities. The embedding medium will slowly de-stain the preparations, however cellular debris within the cells remains clearly visible in form of brown deposits.

### Immunoperoxidase stain for Aβ and Tau protein

For Tau AT8 labelling of human brain we utilized the mouse anti-AT8 antibody (Thermo Fisher Scientific #MN1020) 1:1600 dilution; A*β* 1-42 was detected with the rabbit monoclonal mOC 64 antibody (AbCam # 1ab201060) 1:1000 dilution. Immunostaining was performed in the histology lab at the University of Vermont Medical Center. We labelled 8 µm paraffin embedded brain slices using a Leica Bond-3 autostaining system with heat-based antigen retrieval. A low pH buffer solution was utilized (AR9961; Leica, Bond Epitope Retrieval Solution 1, 10 min). The incubation time for the antibodies was 15 min. Signal detection was achieved using a polymer detection system (DS9800: Leica, Bond Polymer Refine Detection).

### Luxol H&E staining of Autopsy Tissue

This procedure was conducted at the University of Vermont Medical Center in the histology lab. Brains were fixed for the duration of at least one week in 10% neutral buffered formalin at room temperature. The hippocampal tissue was dissected from the brains and dehydrated in a graded series of ethanol. Embedding in paraffin was performed through infiltration with 100% xylene (3x 40 min), prior to embedding in paraffin (25 min). Sections were cut at 5-10 μm thickness and mounted on glass slides.

Slides were rinsed in xylene (2x), 100% and 95% EtOH (1x each) prior to immersion in 0.1% Luxol fast blue solution (Alfa Aesar, A15395) at 60 °C overnight. Excess stain was removed with 95% EtOH. Slides were rinsed in distilled water and placed in a 0.05% lithium carbonate solution (2 min) followed by 70% EtOH (2 min). After additional incubation in 0.05% lithium carbonate (1 min), preparations were rinsed in 70% EtOH (1 min) and distilled water. Slides were then stained with hematoxylin (Leica 3801571) for 1 min. Sections were washed (1 min) in water, immersed in defining solution (1 min, Leica 3803598), and rinsed in water (1 min). To enhance staining quality, preparations were suspended in blue buffer (1 min, Leica 3802918), washed in distilled water (1 min), and 95% EtOH (1 min). Sections were immersed in Eosin (2 min, Leica 3801619) and rinsed in 95% EtOH (1×30 sec and 1x 1 min). For permanent storage, water was removed by rinsing in sections in 100% EtOH and xylene (2×1.5 min each). Sections were permanently mounted using Permaslip Mounting Medium and Liquid Coverslip (Alban Scientific, Inc.).

### RNA-scope *in-situ* RNA hybridization

For the detection of mRNA for human AQP4, glial fibrillary acid protein, pan-quaking, caspase3, Sox10, and myelin basic protein, we carried out in-situ RNA hybridization. Formalin-fixed paraffin embedded human hippocampal sections (5 µm) were freshly sectioned and treated with Bio-Techne designed complementary probes to all tested proteins (Bio-Techne AQP4 # 482448; MBP # 411058-C2; pan-Quaking #416358-C3; GFAP #311808-C2; Caspase3# 436568-C3; Sox10# 484121-C4) at 1x dilutions. Visualization of target-specific signals through the use of tandem scaffolds was achieved with enzyme-linked amplifier probes coupled to both probes allowed. For control purposes, RNA integrity in the tissue sections was tested using in situ hybridization for low, medium, and high level expressed human ‘housekeeping’ genes which allowed for the assessment of overall RNA integrity in the stained sections. We utilized 3-plex positive control probes. Channel C1: POLR2A (RNA polymerase II subunit) moderate-to-low expressor; Channel C2: PPIB (peptidylprolyl isomerase B) moderate-high expressor; Channel C3: UBC (ubiquitin C) highest relative expression level. For negative controls we utilized the bacterial gene dapB. We assessed RNA integrity (PPIB-positive and dapB-negative) for all samples. Visualization of the aquaporin 4 RNA will be achieved via laser scanning confocal microscopy at 40x magnification. The labeling procedure was carried out using a Leica Bond RXm (Leica Biosystems, Deer Park, IL) using protocol ACD Multiplex 2 Plex and all RNAscope 2.5 LSx supporting protocols. Sample preparation consisted of 20 minutes of bake and dewax followed by a 20 min antigen retrieval in epitope retrieval 2 and a 15 min protease treatment at 95°C. AQP4 was labeled with OPAL 570 and GFAP was labeled with OPAL 650. Samples were counter-stained with DAPI.

### Light microscopic image acquisition

Histological brain sections were examined and captured using an Olympus light microscope with digital image acquisition. Figures were created using Adobe Photoshop (RRID:SCR_014199) and Illustrator (RRID:SCR_010279).

### Immunohistochemistry negative controls

The antibodies utilized here have been well-established in the human brain. We have routinely carried out negative controls in which the primary antibodies were omitted to exclude that the observed strong fluorescence may be attributed to autofluorescence in the tissue.

## Results

### Myelinated ependymal tanycytes form swell-bodies and varicose cell processes that project into the hippocampal formation

The ependymal lining in the hippocampal formation of mouse, rat, and humans that borders the ventral horn of the lateral ventricles contains an interconnected network of myelin-forming tanycytes (Fig 1A-O). Investigation of alternating semi- and ultrathin sections through the ependymal lining in rat hippocampus shows that these ependymal macroglia give rise to a dense network of myelinated processes (Fig 1). These slender projections pull into the hippocampal formation and form numerous, characteristic electron-lucent varicosities (Fig 1M, Fig 2). The light- and electron-microscopic appearance of these myelin-forming tanycytes whose somata contain large nuclei and a structured cytoplasm is distinctly different from circular organelles that form along the myelinated processes (Fig 1B insets 4-6, D-T). The latter have electron-lucent, transparent lumina that are void of both cytoplasm and associated organelles except for nucleus-like organelles that are physically associated with myelinated tanycyte processes (Fig 1O-R). Cell processes around the circumference of these circular organelles appear AQP4 immunoreactive (Fig 1Q). Due to different swell patterns in these translucent, circular organelles, we refer to them as ‘swell-bodies’ (see [28] for more detail on swell-bodies). As described in more detail below, swell-bodies form receptacles that emanate from the nucleus-like structures (Fig 1S, T). Due to clear distinctions from true cell nuclei listed in more detail below, we refer to these nucleus-like structures as ‘tanysomes.’ Correlative light- and electron-microscopy of well-preserved rat hippocampus demonstrates that the myelinated tanycyte processes project into the stratum pyramidale and form close contact with neuronal dendrites, somata and axons. Neuronal processes were unambiguously identified based on their physical association with neuronal somata in longitudinal sections (Fig 2). Interestingly, the myelinated tanycyte processes form varicose projections into adjacent cells, including neuronal somata and their associated neurites as demonstrated at both light- and electron-microscopic levels (Fig 2B, D). Some varicose projections contain electron-dense material consistent with the ultrastructural appearance of cellular waste (Fig 2D, E, H).

**Figure 1.**
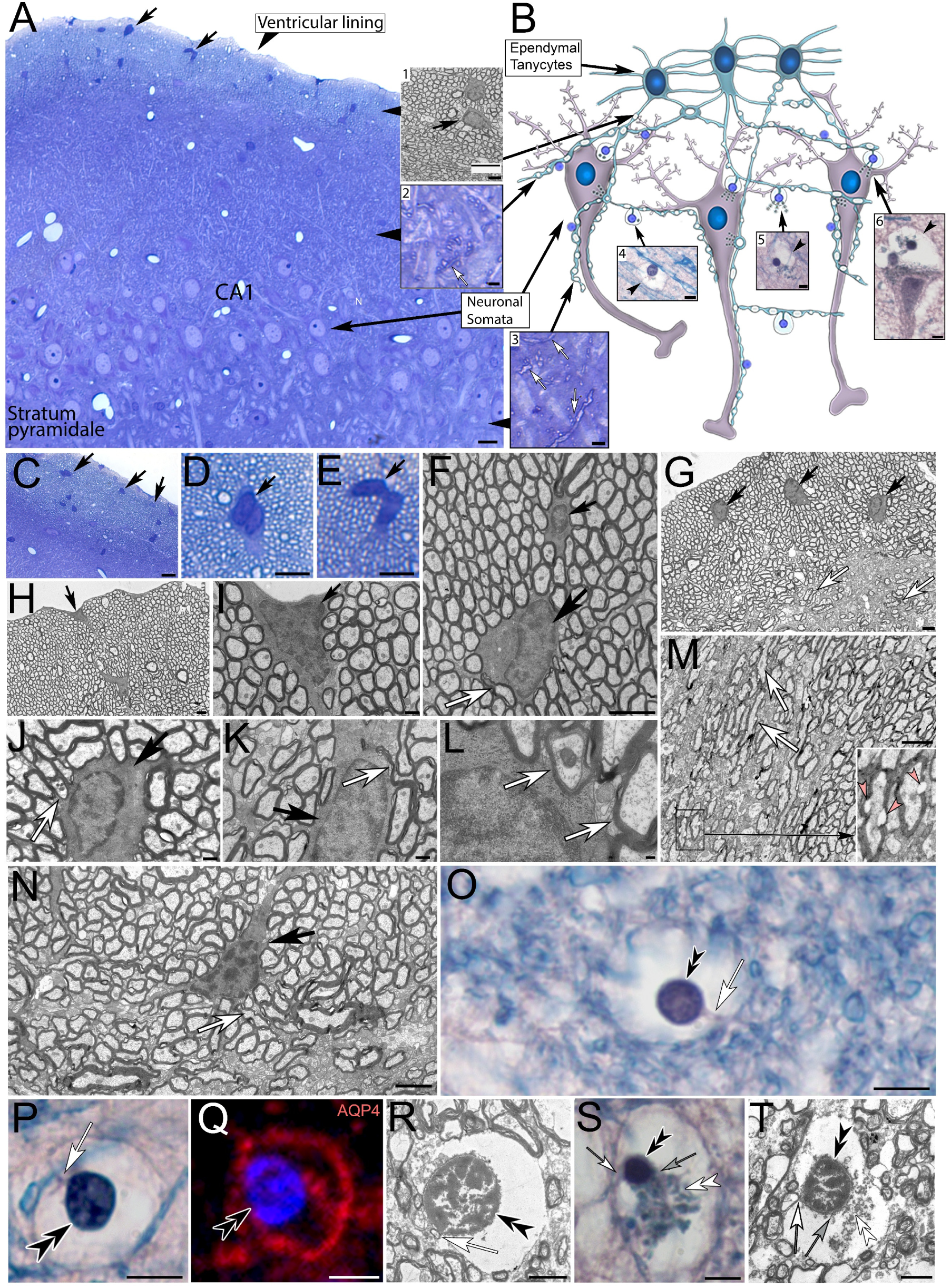
Histological and ultrastructural depiction of ependymal tanycytes in rat (*A-N*) and human (*O-T*) hippocampus. **(A)** Toluidine blue stained 1-µm semithin section of rat hippocampus shows the location of myelin-forming ependymal tanycytes (*black arrows*) in the ventricular lining (*inset 1*). The myelinated tanycyte processes project into the hippocampal formation and form close contact with the somata and neurites of pyramidal cells (*arrow in inset 2*). **(B)** Schematic depiction of the proposed connectivity between the myelinated, ependymal tanycytes and hippocampal pyramidal cells in the CA3, CA2, and CA1 regions. The varicosity-forming slender tanycyte processes project into the stratum pyramidale and form varicose processes along neurons (*arrows in inset 3*). In both rodent and human hippocampus, the varicose tanycyte processes give rise to translucent swell-bodies that contain nucleus-like tanysomes (*arrowheads in insets 4-6*). **(C-E)** Higher magnification of ependymal tanycyte somata in rat hippocampus shows the multinucleated and interconnected nature of this cell network. **(F-N)** Ultrastructural depiction of the tanycytes using alternating semi- and ultrathin serial sections demonstrates well preserved somata with clear nuclei and structured cytoplasm (*black arrows*). The well-defined cytoplasm gives rise to myelinated cell processes (*white arrows*). These slender cell processes project from the ependymal lining into the hippocampal formation (*M, white arrows*) and form characteristic varicosities along their longitudinal axes (*red arrowheads in inset*). **(O-T)** Swell-bodies with their translucent lumina shown in Luxol H&E-stained paraffin sections (*O, P, S*), AQP4 immunolabeling (Q) and at the ultrastructural level (*R, T*). Each swell-body contains one or more small round nucleus-like tanysomes (*black double arrowheads*) that are associated with myelinated tanycyte processes (*white arrows*). The swell-bodies in *S* and *T* form receptacles that emanate from the tanysomes (*white double-arrowheads*). *Grey arrows in S, T:* association between tanysomes and emanating receptacles. **Scale bars:** A: 10 µm (inset1: 2 µm, inset2: 3 µm); B: 5 µm (inset3: 3 µm, insets4-6: 5 µm); C: 10 µm; D, E: 5 µm, F-H :2 µm; I, J: 500 nm; L: 100 nm; M, N: 2 µm; O-Q: 5 µm; R: 2 µm; S: 5 µm; T: 2 µm.

**Figure 2:**
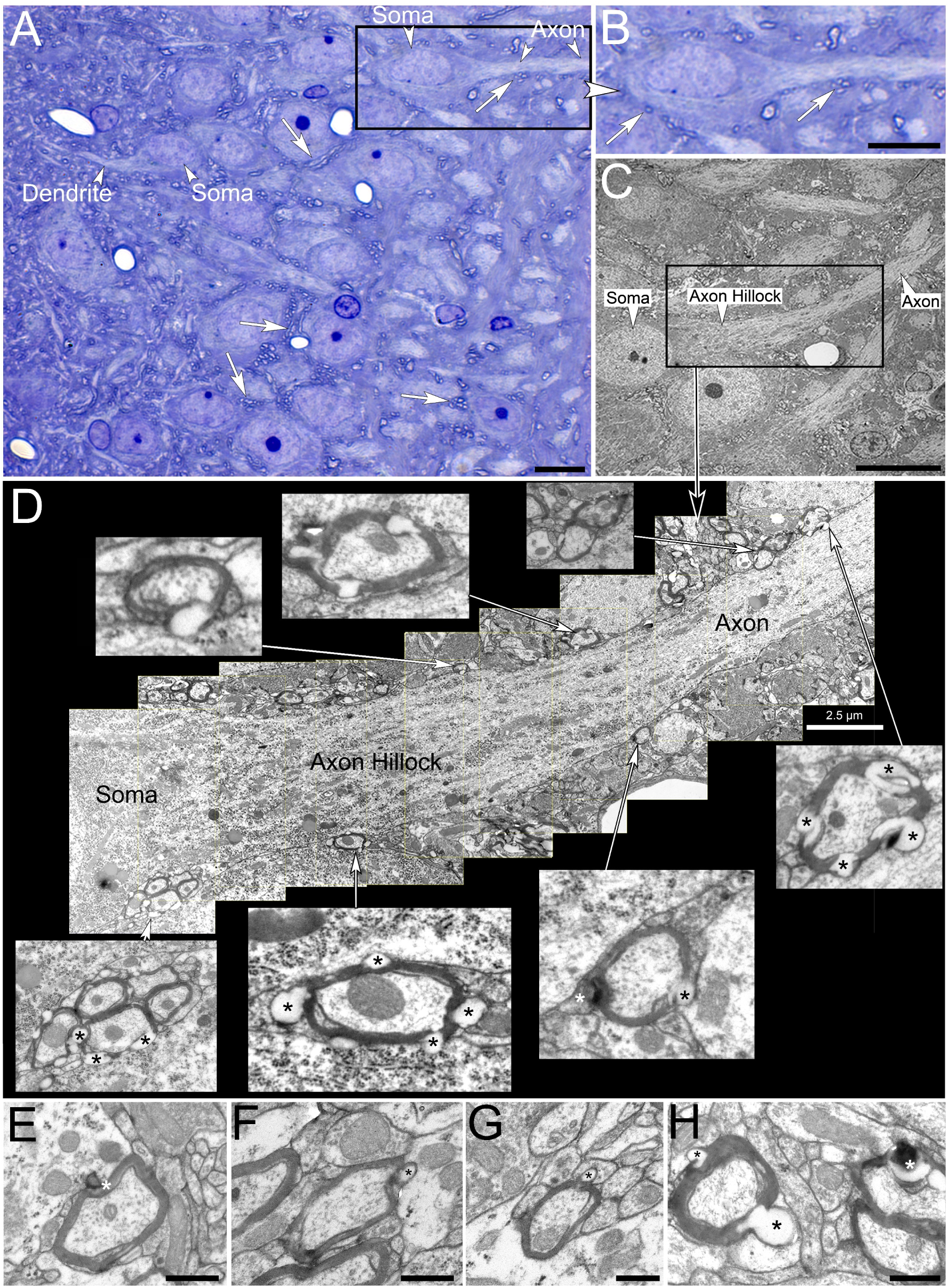
Histological and ultrastructural depiction of connectivity between varicosity-forming myelinated tanycyte profiles and CA2 pyramidal cells in the rat hippocampus. **(A,B)** Toluidine blue stained 1-µm semithin section shows neuronal somata and their dendritic and axonal processes. Numerous myelinated varicosity-forming tanycyte processes (*representative examples indicated by white arrows*) form close contact with the pyramidal cells (*Dendrite, Soma, Axon*). **(C, D)** Representative longitudinal section through the soma, axon hillock and axon of a pyramidal neuron (*C, lower magnification*) shown at higher magnification in *D*. The image montage (*D)* shows the close association between the neuronal axon and varicosity-forming tanycyte processes that are either electron-lucent (*insets in D black asterisks*) or contain electron-dense material consistent with the ultrastructural appearance of cellular waste (*insets in D, white asterisks*). **(E-H)** Representative electron micrographs of myelinated tanycyte processes and their protruding varicosities that project into surrounding cell profiles (*black and white asterisks*). **Scale bars:** A-C: 10 µm; D: 2.5; E-H: 500 nm.

### Swell-bodies in the human hippocampus are associated with myelinated cell processes

Investigation of Luxol H&E-stained human brain sections shows similar varicose, myelinated cell processes that project from the alveus into the hippocampal formation as indicated by Luxol-blue, a histological stain for myelin (Fig 3). In both human and mouse hippocampus, we also observed AQP4-immunolabeled ependymal cells with long varicose slender processes that project into the brain parenchyma (Fig 3C, D, I-K, see also below). As demonstrated in Fig 2 at both light- and electron-microscopic levels such varicose protrusions are absent along neuronal processes that are clearly identified as such based on their association with the neuronal somata visualized in longitudinal sections through neurons. Interestingly, in both Luxol H&E-stained human hippocampus and myelin/AQP4 double labeled mouse hippocampus these processes show a ring pattern whereby strongly Luxol blue-stained (Fig 3H) and myelin-immunolabeled rings are observed in regular intervals along the cell processes reminiscent of stabilizing cartilaginous rings in trachea (Fig 3 H-K; see discussion). As demonstrated in mouse brain AQP4-immunoreactivity is observed in the same cell processes that label for myelin (Fig 3I-K). Whereas the AQP4-signal predominantly labeled the outer circumference of these cell processes, the myelin signal was often associated with circular structures that line the inside of these glial processes.

**Fig 3.**
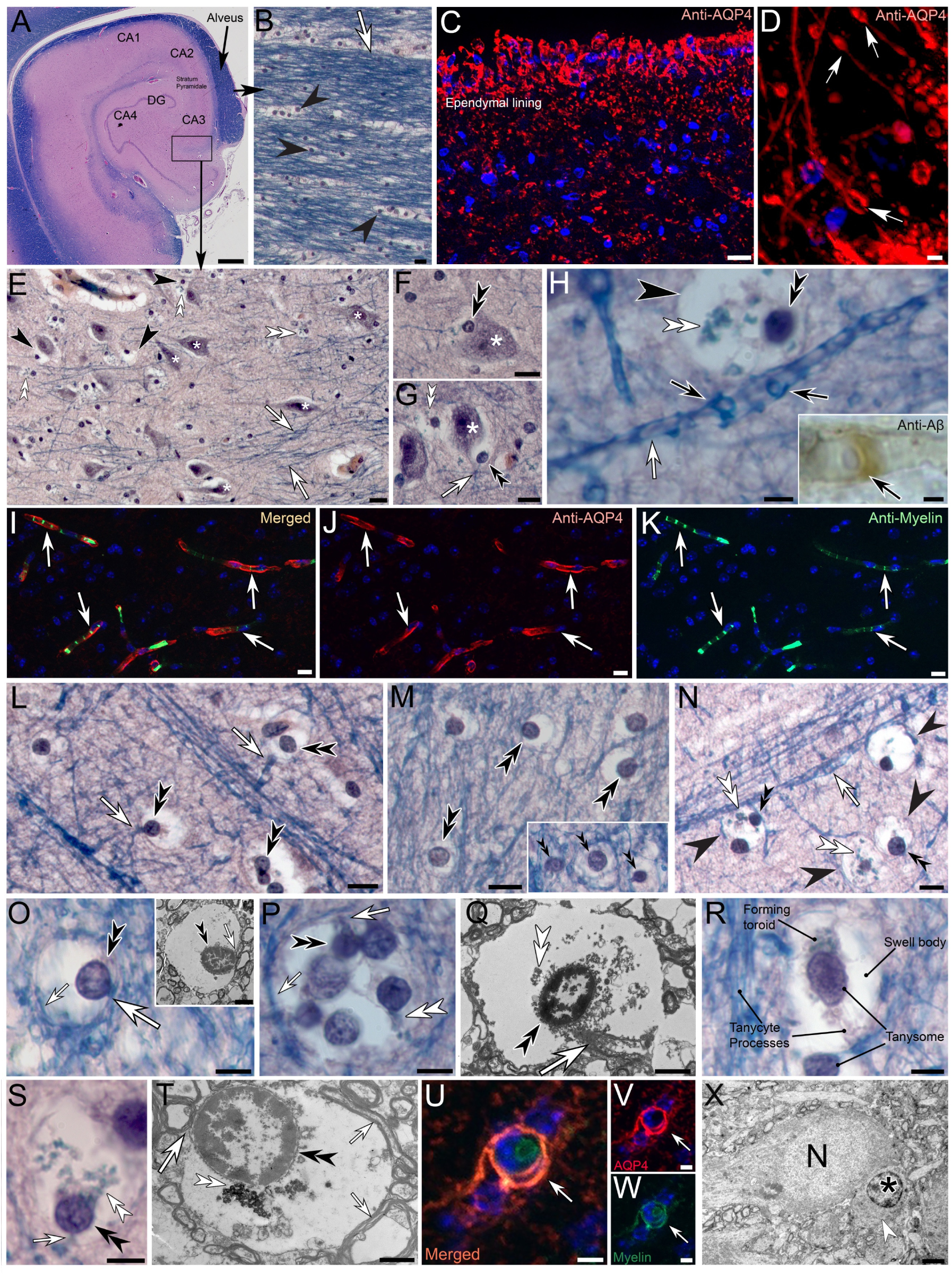
Myelinated Ependymal tanycytes in human and mouse hippocampus project into the hippocampal stratum pyramidale and give rise to waste-internalizing receptacles. **(A-B)** The alveus of human hippocampus contains numerous Luxol H&E-stained ependymal tanycytes that send vast numbers of slender myelinated processes (*white arrow in B*) into the stratum pyramidale of the hippocampal formation. Numerous electron-lucent ‘swell-bodies’ with associated nucleus-like tanysomes are forming along individual myelinated processes (*black arrowheads in B*). **(C, D)** Aquaporin4-immunolabeling of human alveus shows strong immunoreactivity of ependymal tanycytes (C) and their long slender processes (D). Higher magnification *(D)* shows varicose protrusions that from along the slender tanycyte processes (*white arrows in D*). **(E-G)** Luxol-blue stained human tanycyte processes (*white arrows*) project into the stratum pyramidale where they form translucent swell-bodies (black arrowheads) with associated nucleus like tanysomes (*black double arrowheads*). Please note the close association of swell-bodies with neuronal somata (*asterisks*). **(H)** Tanycyte (*white arrow*) with associated translucent swell-body (*black arrowhead*) with a receptacle forming (*white double arrowhead*) tanysome (*black double arrowhead*). Swelling ring structures (*black arrows*) that line the inside of swelling tanycyte processes are visible in Luxol H&E-stained and Aβ-immunolabeled (*black arrow in inset*) tanycyte processes. **(I-K)** AQP4 (*red*) and myelin (*green*) immunolabeling of mouse hippocampus shows double labeled tanycyte processes that contain myelin-immunoreactive circular profiles that line the inner lumina of the cell processes (*white arrows*) consistent with our observations in human brain (*H*). **(L-N)** Tanycyte processes (*white arrows*) with associated swell-bodies (*black arrowheads*) in human hippocampus. *White double arrowheads:* forming receptacles; *Black double arrowheads:* Tanysomes. *Inset in M*: Forming tanysomes. **(O-T)** Structural characteristics of swell-bodies, associated tanysomes (*black double arrowheads*) and receptacles (*white double arrowheads*) in human hippocampus. Both light- and electron-microscopic images show the electron-lucent nature of the swell-bodies that contain between one and 6 tanysomes shown in Luxol H&E-stained preparations. The association of tanysomes with myelinated tanycyte processes is clearly visible (*white arrows*). Tanysomes give rise to toroids and receptacles some of which appear electron-dense at the ultrastructural level (*white double arrowheads in T*). Please note the myelinated tanycyte profiles that line the circumference of swell-bodies (*small white arrows in O, P, T*). **(U-W)** AQP4/Myelin immunolabeled swell-body in mouse hippocampus shows the double-labeled nature of myelinated AQP4-immunoreactive tanycyte profiles (*white arrow*) around the periphery of swell-bodies. **(X)** Typical structured appearance of a glial nucleus (*asterisk)* and surrounding cytoplasm (*white arrowhead*). **Scale bars:** A: 500 µm, B:10 µm, C: 20 µm; D: 5 µm; E-G: 10 µm, H: 5 µm, Inset 3 µm; I-N: 10 µm; M: 20 µm; N: 10 µm; O: 5 µm, (*Inset*) 2 µm; P: 5 µm Q 2 µm R: 5 µm; S: 5 µm; T: 2 µm; U-W: 5 µm; X: 2 µm.

Both the AQP4-immunoreactive and translucent nature of swell-bodies that form along these myelinated processes is consistent with the hypothesis, of an AQP4-mediated swelling of these organelles (see discussion). Swell-bodies are clearly visible in Luxol H&E-stained human hippocampus (Figs 3, 4) and AQP4/myelin immunolabeled mouse hippocampus (Figs 3 U-W; 4 P-R). In human hippocampus each swell-body contains between one and six tanysomes (Fig 3L-T). Unlike typical cell nuclei, these organelles give rise to receptacles (Figs 3T, 4A-D; see also below) that often appear electron-dense at the electron-microscopic level consistent with the ultrastructural appearance of cellular waste (Fig 3T *white double arrowhead*). Tanysomes have three characteristics that clearly distinguish them from conventional cell nuclei.

**Fig 4.**
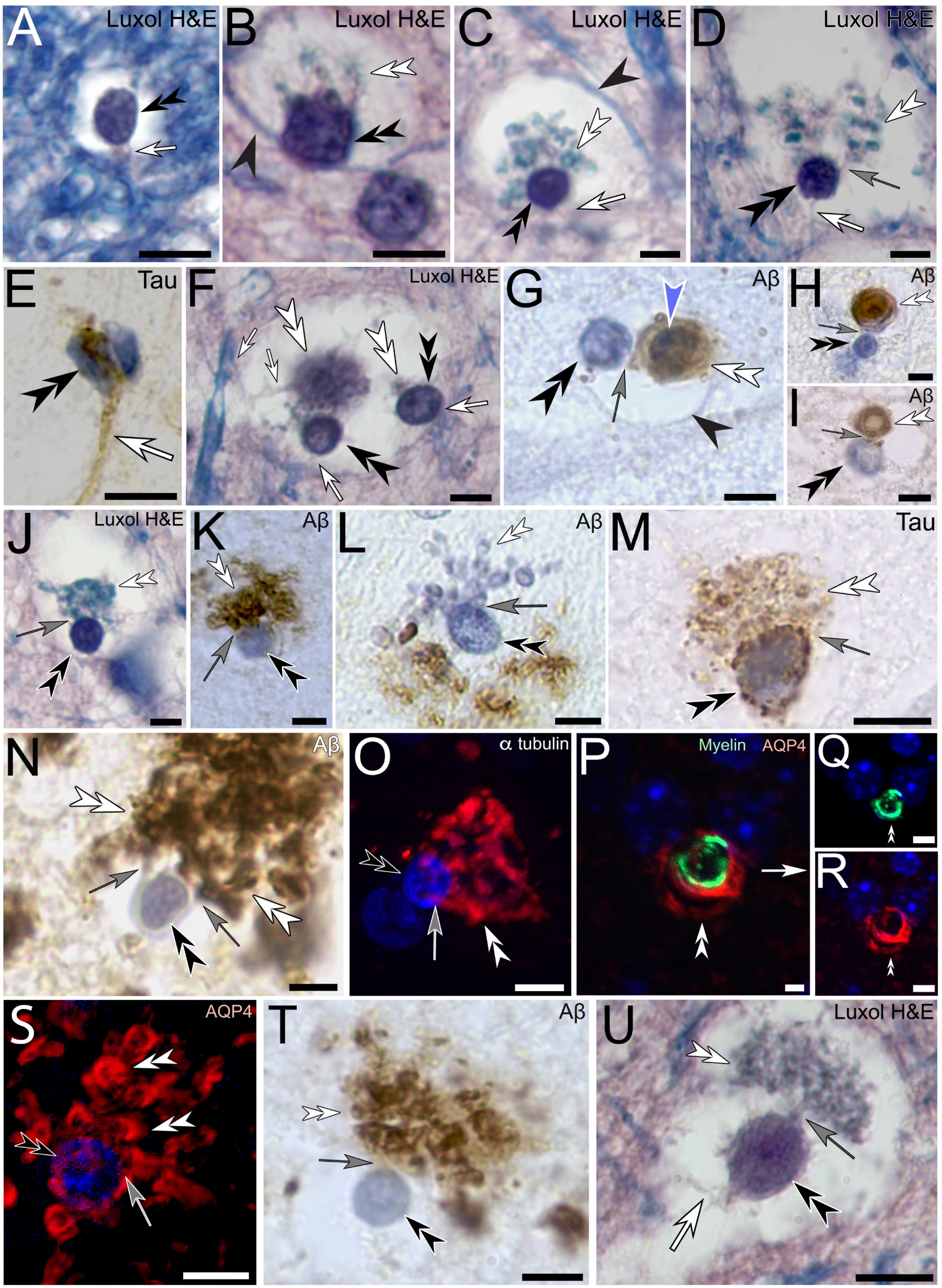
Tanysomes in human and mouse hippocampus give rise to toroids and receptacles that are immunoreactive for Aβ, hyperphosphorylated tau protein, myelin, AQP4 and α-tubulin. **(A-O)** Human hippocampal tanysomes within swell-bodies give rise to a network of membranous receptacles and toroids (*white double arrowheads*). The physical attachment between tanysomes (*black double arrowheads*) and tanycyte processes (*white arrows*) and receptacles (*grey arrows*) is clearly visible especially in panel E. Nuclear stain is observed in tanysomes and although weaker in emanating toroid-shaped structures (*blue arrowhead in G*) and receptacles (*L*). The tanysome-derived membrane network in human brain is immunoreactive for Aβ (G, *H, I, K, L N,T*), Tau (*E, M*), α-tubulin (*O), and* AQP4 (*S*). **(P-R)** Immunolabeling of mouse hippocampus for anti-AQP4 (*red*) and myelin (*green*) demonstrates that toroids that emerge from tanysomes are immunoreactive for both epitopes. **(S-U)** Tanysome-derived receptacles in preparations immunolabeled for anti-AQP4 (S), anti-Aβ (T) and stained for Luxol H&E (U) consistently demonstrate their physical association with tanysomes (*grey arrows*). Scale bars: A-U: 5 µm.

### Tanysome versus conventional cell nucleus

Despite the positive nuclear stain, the following features distinguish tanysomes from conventional nuclei found in cell bodies in both non-AD-affected and AD-affected human hippocampus: *(i)* the emergence of the aforementioned receptacles from these organelles (Fig 3 S, T; Fig 4 see more detail below), *(ii)* our observation that tanysomes are physically connected with myelinated tanycyte processes (Figs 1O, P, R, T; 3 O, Q, R, T; 4). This physical association is particularly visible in anti-tau immunolabeled AD-affected hippocampus (Fig 4E) and *(iii)* the emergence of swelling toroid-shaped structures from tanysomes (Fig 4G-I). In contrast to the associated tanysomes from which they emerge, these toroids show immunolabeling for Aβ demonstrating that these are not dividing cell profiles. In AD-affected brain these toroids swell to widely varying diameters of >20 μm and give rise to numerous receptacles (see below). Interestingly, close examination shows that both receptacles and toroids that emanate from tanysomes show nuclear stain that is often weaker compared to the nuclear stain intensity in associated tanysomes. Clear examples are shown in Figs 4G, L. This weak nuclear stain is often hard to detect since nuclear stains show only weak interactions with mRNA compared to DNA.

### Histology and histopathology of swell-bodies

The lumina of swell-bodies that are void of receptacles consistently appear translucent at the light microscopic level in Luxol H&E-stained preparations and electron-lucent at the electron-microscopic level (Figs 1 O,P,R; 3 O and inset; 4 A) unlike the structured cytoplasmic appearance in neuronal and glial cell somata (Fig 3E, F, G, X). In other swell-bodies tanysomes give rise to protrusions that form either receptacles (Fig 4B-D) or toroids (Fig 4F-J). In AD-affected brain tissue these receptacles proliferate, swell, and protrude out of the swell-body lumina (Fig 4M, N, S, T). Immunolabeling demonstrates that these receptacles stain for Aβ, α-tubulin and AQP4 (Fig 4). The appearance and diameters of these interconnected receptacles vary widely from electron-lucent with diameters of ∼20 nm to electron-dense with diameters of up to 1 μm.

### Tanycyte-derived receptacles project into neuronal somata

Immunolabeling demonstrates the AQP4-IR nature of waste receptacles that emanate from tanysomes (Figs 4S; 5A). Investigation of Luxol H&E-stained brain sections visualize the strands of receptacles that emanate from tanysomes (Fig 5B). In both AD-affected and AD-unaffected human hippocampus swell-bodies are observed contacting neuronal somata and forming receptacles that project into neuronal somata (Fig 5C-G). Please note the close association of the swell-body with the neuronal somata in non-AD-affected tissue, which leads to a convex indentation consistent with hydrostatic pressure exerted on the soma (Fig 5G). In AD-affected brain tissue, the receptacles appear hypertrophic compared to non-AD-affected tissue (Figs 5C-F, 6A compared to 5G, 6B). These findings are consistent with ultrastructural observations that show numerous waste receptacles that have the ultrastructural appearance of lipofuscin within neuronal somata (Fig 5I). The dense obstruction of AD-affected neuronal somata with waste receptacles is apparent at both the light- and electron microscopic levels (Figs 5D, E; 6A light microscopy *white double arrowheads; Fig 5H, J electron microscopy*) and is often associated with cytoplasmic depletion (Fig 5F, J). In contrast, receptacles in non-AD-affected human hippocampus are less prominent (Fig 5G, I; 6B, C). Interestingly, the intraneuronal receptacles consistently show immunolabeling for Aβ in healthy neurons (Fig 6B). Correlative semithin and ultrathin sectioning were performed on healthy neurons with intracellular receptacles that demonstrate the waste-internalizing nature of these receptacles consistent with the appearance of waste-containing lipofuscin (Fig 6C, D; see discussion). Swell-bodies that form extracellular receptacles in AD-affected tissue proliferate and swell to large diameters resembling structures known as Aβ plaques (Fig 6E-I). The emergence of these receptacles from tanysomes is clearly visible as indicated by the grey arrows in the provided images. Immunolabeling for hyperphosphorylated tau protein in AD affected human brain demonstrates that this protein is also associated with receptacles that emanate from tanysomes that swell in AD-affected brain (Fig 6 J-M).

**Fig 5.**
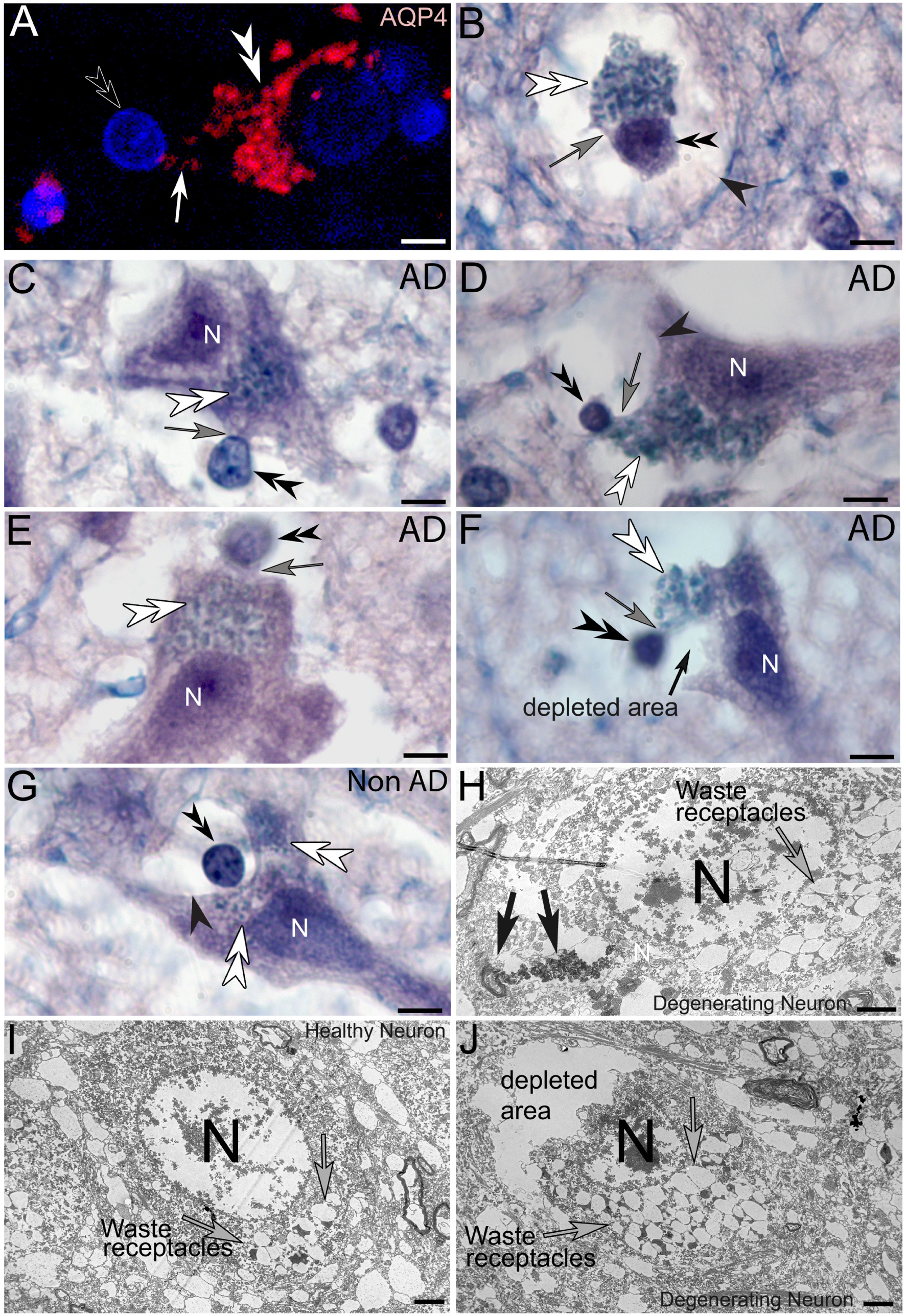
Tanysome-derived waste receptacles in human hippocampus project into neuronal somata. **(A)** Tanysome-derived waste receptacles are immuno-positive for aquaporin-4 (*white double arrowhead*). **(B-G)** Luxol H&E-stained preparations show the characteristically blue stained tanysome-derived waste receptacles that differentiate in swell-bodies (*panel B*) project into pyramidal cell somata. *Black double arrowhead*s: Tanysomes; *White double arrowhead*s: Waste receptacles; *Black arrowheads*: Outer margins of swell-bodies; *N*: Neuronal somata. **(H-J)** Ultrastructural depiction of healthy and degenerating neuronal somata demonstrates the abundance of swelling waste receptacles and gradual depletion of cytoplasmic areas in degenerating somata (*H, J*) compared to the much sparser number of waste receptacles in a healthy neuron that shows an intact cytoplasmic structure (*I*). Please note the waste accumulation associated with a waste receptacle strand that emanates from a myelinated cell profile (arrows in H) consistent with strands of waste receptacles observed at the light microscopic level (C-G). Scale bars: A-G: 5 µm; H-J: 2 µm.

**Fig 6.**
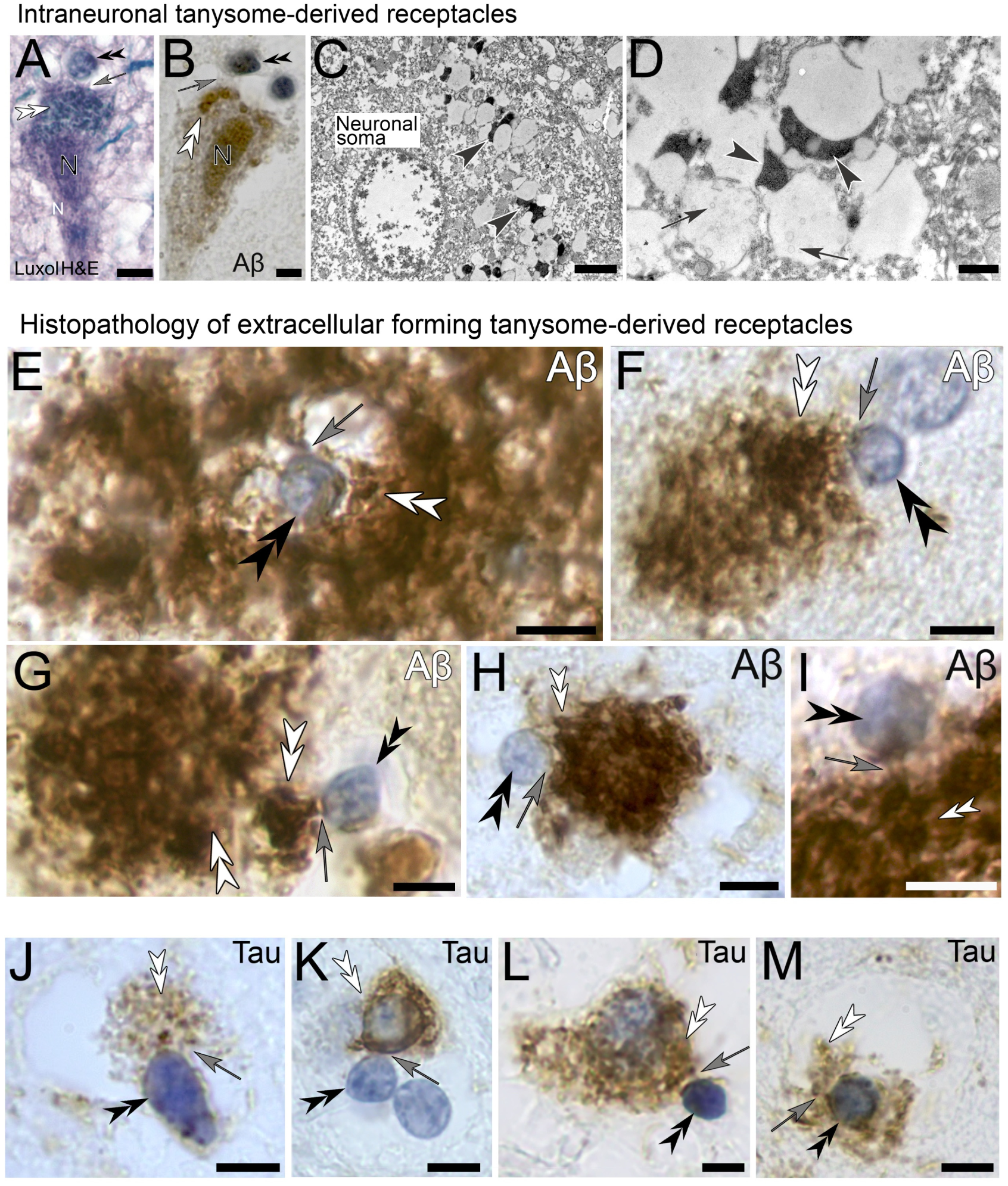
Amyloid beta and tau immunolabeling of tanycyte derived waste receptacles in human hippocampus. **(A, B)** Tanysomes form receptacles that project into AD-affected (*A, Luxol H&E stain*) and non-AD-affected (*B, Aβ labelled*) neurons. The observed intraneuronal Aβ immunoreactivity of the receptacles is consistent with observations of extracellular receptacles. **(C, D)** Ultrastructural depiction of waste receptacles in a healthy neuron that showed moderate amounts of waste receptacles at the light microscopic level. The receptacles consist of circular, electron-lucent components that contain vesicular structures (*black arrows*) and associated electron-dense components (*black arrowheads*). The latter are consistent with the appearance of cellular waste at the electron-microscopic level. **(E-I)** Amyloidβ immunoreactive hypertrophic receptacles emanate from tanysomes in AD-affected brain samples forming Aβ plaques. (J-M) Tau-immunoreactive receptacles that emanate from tanysomes in AD-affected brain tissue. *Black double arrowhead*s: Tanysomes; *White double arrowhead*s: Waste receptacles; *Grey arrows:* Association of emanating receptacles from tanysomes. Scale bars: A, B: 5 µm; C: 2 µm; D: 500 nm; E-M 5 µm.

### Protein and mRNA expression in tanysomes and associated toroids and receptacles within swell-bodies

Interestingly, both toroids and receptacles that emerge from tanysomes often appear weakly stained for nuclear stain (Fig 7B, D). To test our hypothesis that swell-bodies are discrete ‘waste-disposal’ units throughout the hippocampal formation that are ‘ready’ to quickly differentiate functional waste-internalizing receptacles in response to external cues we have investigated mRNA expression and correlative immunolabeling for the translated proteins. Based on our previous findings in spider, mouse, rat, and human brain (Fabian-Fine et al. 2024) we postulated that required proteins are (*i*) mRNA-binding proteins of the quaking family to anchor required mRNA within the toroids, (*ii*) myelin basic protein and glial acid fibrillary protein (GFAP) that likely provide the structural basis for these receptacles, (*iii*) AQP4 that likely plays a functional role in the formation of a convective flow toward the receptacles, (*iv*) Caspase 3 that likely catabolizes cellular waste, (*v*) Sox 10 that governs differentiation, and (*vi*) presenilin 1 and APP, both proteins contribute to the formation of amyloid beta which we postulate provides structural support for the forming receptacles to prevent the collapse of waste-receptacles during AQP4-mediated waste intake (see discussion). As demonstrated in Figs 7 and 8 we detected mRNA for all tested probes within swell-bodies. GFAP was demonstrated in a previous publication, however we have included a representative image into summary Fig 12 to demonstrate the robustness of these experiments. To test that the expressed proteins are indeed present within swell-bodies and associated receptacles we have furthermore conducted extensive immunolabeling. Of particular interest is the observed AQP4-immunolabeling of swell-bodies and emanating receptacles in AD-affected brain tissue (Fig 7I, J, L, M) this labeling is consistent with the observed mRNA expression in swell bodies (Fig 7 E4, G4). Please note the clear distinction between the unlabeled clear lumina characteristic for swell-bodies (Fig 7 H, K, N). In contrast, AQP4-immunoreactive cells show strong immunolabeling within their somata consistent with immunolabeled cells (Fig 7O, inset). Immunolabeling for Aβ (Fig 7, 8), presenilin 1, APP and caspase3 (Fig 8) were also consistent with our observed gene-expression patterns. Immunolabeling of non-AD-affected brain demonstrates that the tanysome-derived receptacles label for Aβ, however the labeling intensity is weaker compared to AD-affected tissue (Fig 8W, X). In non-AD affected brain swell-bodies and associated receptacles and toroids mostly lack excessive proliferation of Aβ-labelled receptacles (for quantitative analysis please see [28]). Negative controls showed only weak background fluorescence in all channels (Fig 8A inset mRNA probe). Positive controls showed the expected gene-expression patterns for the tested housekeeping genes (see Methods) in our RNAscope experiments.

**Fig 7.**
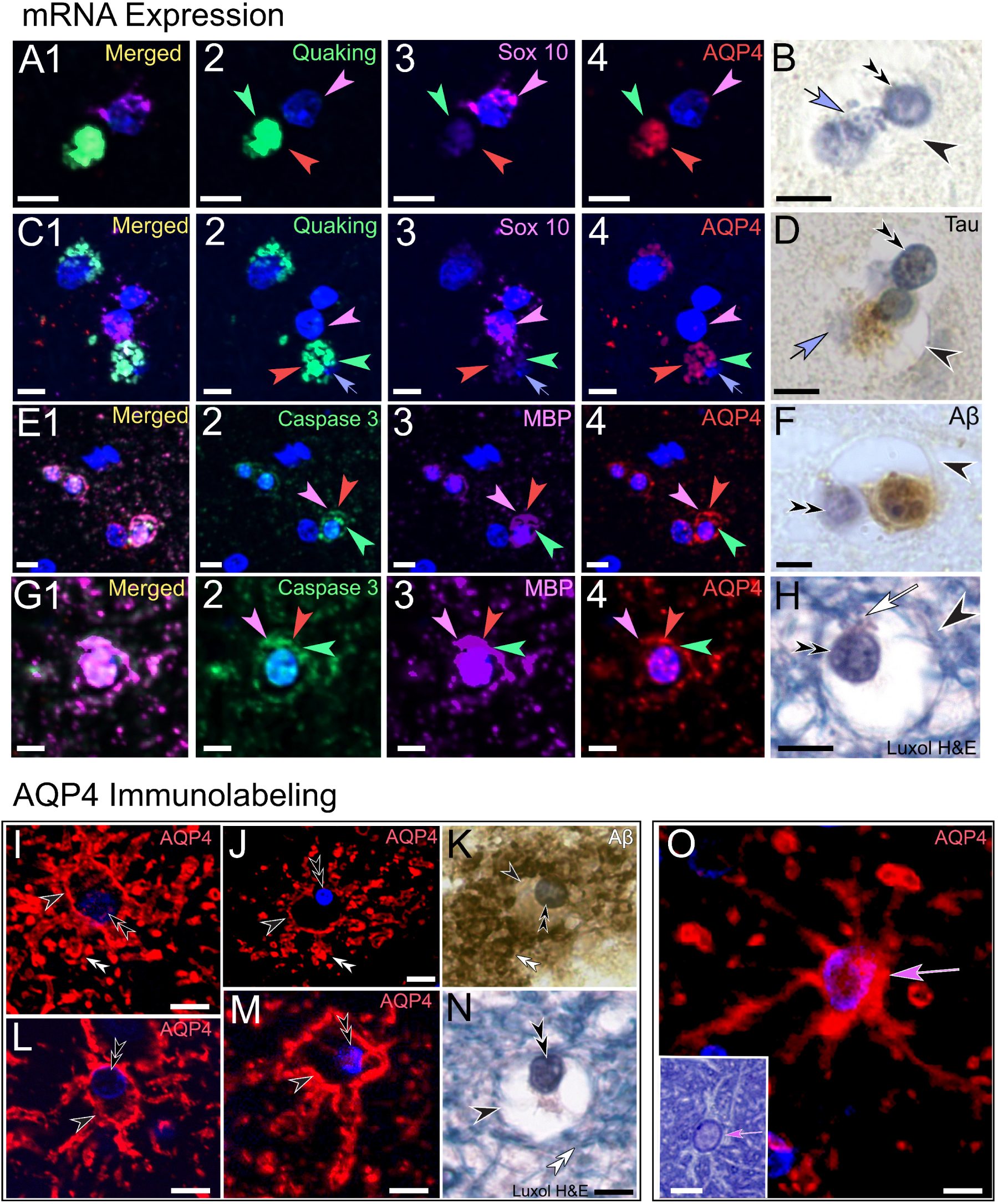
mRNA expression in healthy human hippocampus. **(A1-4)** Swell body with emerging toroid shows mRNA for pan-Quaking (*green arrowhead*) and AQP4 (*red arrowhead*). Sox 10 (*magenta arrowhead*) is restricted to the associated tanysome. **(B)** Histological equivalent shows a swell-body (*black arrowhead*) that contains a tanysome (*black double arrowhead*) that gives rise to a nuclear-stained receptacle-containing toroid (*purple arrow*). **(C1-4)** Receptacle-forming swell-body shows pan-quaking (green arrowhead) and AQP4-mRNA (*red arrowhead*) associated with the forming receptacles, whereas Sox10 is associated with the tanysome (*magenta arrowhead*). **(D)** Amyloidβ immunolabeled equivalent of a swell-body in a similar stage of receptacle formation. Please note the subtle nuclear stain of the receptacles that are not stained for Aβ (*purple arrow*). **(E1-4F, G1-4, H)** Swell-bodies labeled for caspase 3 (*green arrowheads*), myelin basic protein (*MBP, magenta arrowheads*) and AQP4 (*red arrowheads*) show co-localization of all three mRNAs within the swell-bodies. Histological equivalents are shown in *F (Aβ-immunolabeled*) and *H (Luxol H&E stain*). *White arrow in H*: association of the tanysome with the tanycyte. **(I-N)** Comparison of immunolabeled (*I, J, L, M: AQP4, K: Aβ*) and Luxol-H&E (*N*) stained AD-affected human well-bodies (*black arrowheads*) shows their consistently clear and unstained lumina. Numerous receptacles can be observed around the periphery of the swell-bodies (*white double arrowheads*). *Black double arrowheads*: Tanysomes. **(O)** In contrast, an immunolabeled cell profile in AD-affected brain with a clearly AQP4 immunolabeled soma (*pink arrow*) and associated slender processes resembling the typical shape observed in histological depictions of astrocytes (*pink arrow in inset*). Scale bars: 5 µm.

**Fig 8.**
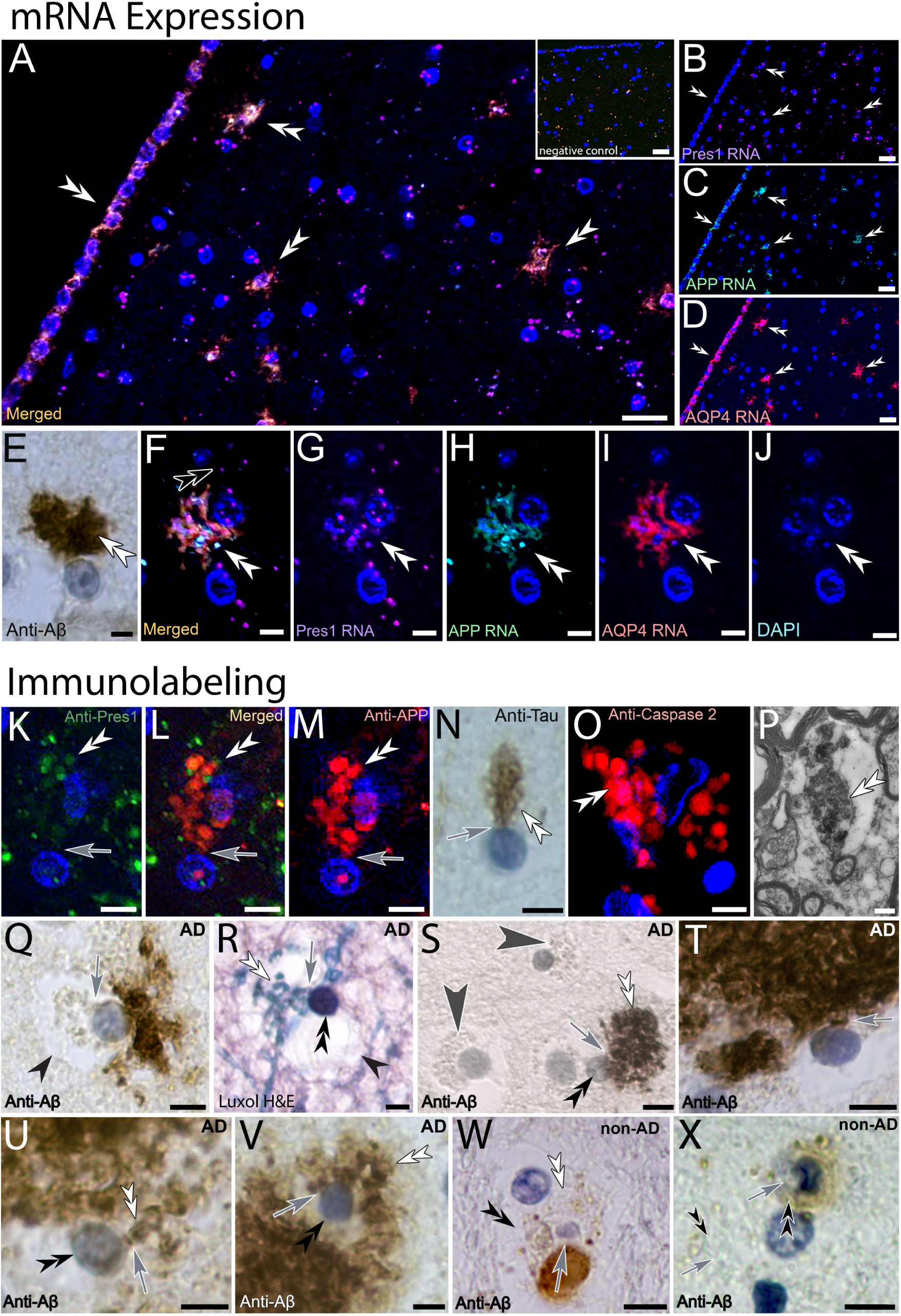
Amyloid β-related gene and protein expression in AD-affected human hippocampus. **(A-D, F-J, L-P)** Ependymal tanycytes in the ventricular lining show RNA-expression for Presenilin1 (*Pres-1*, *magenta*), Amyloid Precursor Protein (*APP*, *green*) and Aquaporin 4 (*AQP4*, *red*). Higher magnification (*F-J, L-P*) shows that gene expression is associated with tanysome-derived toroids and receptacles (*DAPI nuclear stain*). **(E)** Tanysome-associated waste receptacles are strongly Aβ**-** immunoreactive (*white double arrowheads*) consistent with observed Pres1 and APP gene expression shown in *F-J*. **(K-O)** Tanysome-associated receptacles show immunolabeling for Pres1 (green K, L), APP (red, L, M), Tau protein (N) and Caspase 2 (O). (P) The appearance of the immunolabeled receptacles is consistent with the ultrastructural appearance of receptacles that appear electron dense. (Q-X) Depictions of AD-affected swell-bodies that show the emergence of receptacles from swell-body associated tanysomes (Q, S, T, U, V: Ab immunolabeling; R: Luxol H&E stain; W: Tau-immunolabeling; X: anti-α tubulin immunolabeling) consistent with the observed mRNA expression for Pres1, APP both of which are important for the formation of Aβ. **Scale bars:** A-D: 20 μm; E-O: 5 μm; P 500 nm; Q-X: 5 µm.

### Functional evidence for AQP4-mediated waste intake into ependymal tanycytes

To further test our hypothesis that the observed electron-dense material observed in tanycyte-derived waste receptacles is internalized cellular waste, we have investigated fluorochrome uptake by ependymal glial cells in living mouse hippocampus. This preliminary experiment was designed to test the general validity of our experiments. If we postulate that ependymal tanycytes, swell-bodies and associated receptacles internalize cellular waste from extracellular spaces we would expect that this cell network internalizes Cy3 goat anti-rabbit fluorochrome. This goat fluorochrome was chosen because it allowed us to utilize a secondary Alexa 488-coupled donkey anti-goat antibody after fixation of the tested brain to confirm that the internalized fluorochrome was indeed the applied goat-coupled fluorochrome. Control brains were submerged in physiological mouse culture medium only without fluorochrome. The application time of 30 min was chosen to avoid fluorochrome clearance from tanycytes. Within this tested timeframe, the above-described tanycytes and associated ring structures with emanating receptacles appeared brightly fluorescent (Fig 9). In contrast, control preparations lacked fluorescence. These findings are consistent with our hypothesis that the fluorochrome may be internalized by these ependymal tanycytes (Fig 9). The ventricular lining showed a fluorescing, interconnected network of cell processes and canals each containing two small-diameter fluorescent channels within their lumina (Fig 9D insets). These canals project toward the adjacent ventricular lumen (Fig 9E, bottom inset). Immunolabeling of these brain slices with an Alexa 488-coupled donkey-anti goat antibody shows the double-labeled nature of fluorescent structures (Fig 9A-C) consistent with our postulation that the fluorescence observed within the fluorescent cell profiles shows uptake of the Cy3 conjugated antibody from extracellular spaces. Interestingly, tanycytes that border directly onto the ventricle form apical canals that project into the ventricular space (Fig 9A). To conduct preliminary experiments that investigate our hypothesis that the observed internalization of the applied fluorochrome is AQP4-mediated, we have conducted additional experiments in which we have separated and sectioned the two hemispheres of mouse brains. The brain sections of one hemisphere served as control and were kept in culture medium only, the experimental sections from the second hemisphere were immersed in culture medium that contained 60 μM of the selective AQP4-blocker TGN-020. Both control and experimental hemispheres were exposed to the Cy3-conjugated antibody. Analysis of these preparations revealed a significantly reduced fluorescent signal within TGN-020-exposed brain cells compared to the controls in comparable brain areas (Fig 9E-F). Comparisons of Log_10_ mean cellular object pixel intensities demonstrate significantly higher pixel intensity in control (0 µM; −1.888) compared to AQP4-blocked (60 µM TGN-020; −2.278) conditions (Fig 9G). Fluorescence was absent in nearby neurons. We postulate that the observed fluorescence in tanycytes may originate from tanycyte-derived, strongly fluorescent receptacles that internalized waste from extracellular spaces. As demonstrated in Fig 9I, the fluorescent structures strongly resemble in shape and size the receptacles visualized with Luxol blue H&E-stain, Aβ, and AQP4/myelin immunolabeling in both human and mouse hippocampus.

**Fig 9.**
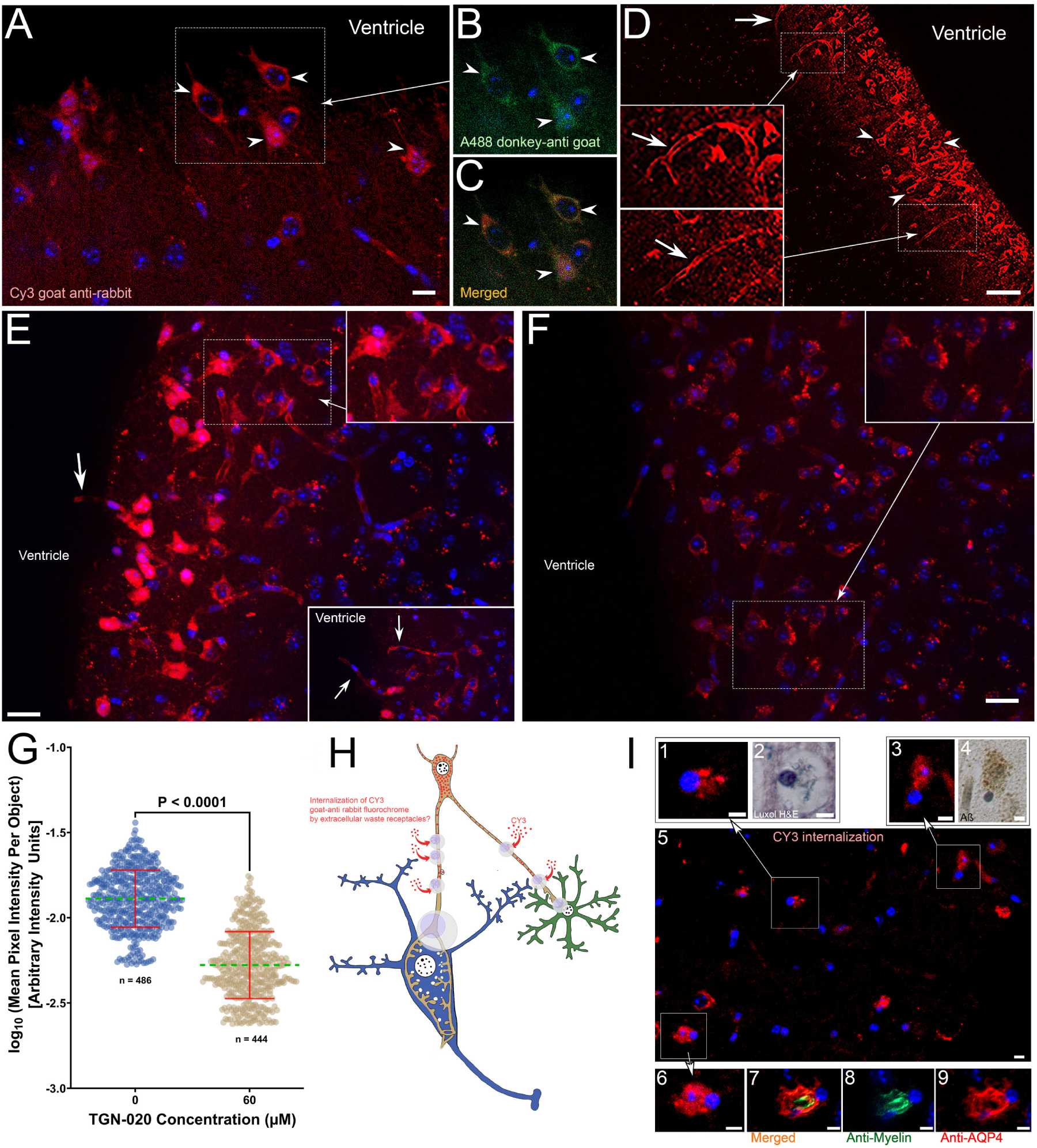
Evidence for aquaporin-4-dependent Cy3 goat anti-rabbit antibody internalization into living mouse hippocampal ependymal tanycytes. **(A-C)** Living ependymal tanycytes that were exposed to Cy3 goat anti rabbit antibody appear fluorescent after 30 min exposure (*white arrowheads in A*). Immunolabeling against goat-protein using an Alexa 488-coupled donkey-anti-goat antibody demonstrates co-localization of both Cy3 and Alexa 488 fluorochromes (*white arrowheads in B, C*) indicative that the red fluorescence within tanycytes originates from internalized Cy3 goat anti rabbit antibody. **(D)** Ependymal tanycytes in mouse alveus show red fluorescence after 30 min exposure of living brain tissue to Cy3 goat anti-rabbit antibody (*white arrowheads*). A fine network of fluorescing processes (*grey arrowhead*) is visible in the stratum pyramidale. *Insets:* Canal structures each containing two fluorescing channels (arrows) are frequently observed projecting into the adjacent ventricle. **(E, F)** Normalized intensity of internalized Cy3 fluorochrome comparing control (0; *E*) and AQP4-blocked (60 µM TGN-020; *F*) conditions. Control cells demonstrate a higher pixel intensity, especially towards the alveus. **(G)** Log_10_ mean cellular object intensity is significantly larger in control (0 µM; −1.888) compared to AQP4-blocked (60 µM TGN-020; −2.278; *t* (928) = 32.666, *p* = 2.44 × 10^−156^, unpaired two-tailed *t*-test). Inset in E: Fluorescent canal structures extending into the ventricle (white arrows). **(H)** Schematic depiction of the proposed waste internalization demonstrated here. We postulate that living tanycytes internalize surrounding fluorochromes via extracellular waste receptacles that are formed within swell-bodies explaining the observed fluorescence in ependymal tanycytes. To visualize cells, Hoechst blue nuclear stain was applied after fixation of the brains. **(I)** Single 3-µm optical confocal section of mouse brain with internalized fluorochromes. Note the strong resemblance of brightly fluorescent tanysome-like areas (*1, 3, 5, 6*) with toroid- and receptacle-forming human tanysomes (*2, 4*) and AQP4/myelin double-labeled mouse tanysomes (*7-9*). Scale bars: A-C: 10 µm, D: 50 µm; E, F: 20 µm; H: not drawn to scale; I: 5 µm.

### Comparison of non-AD-affected and AD-affected human brain is indicative of hypertrophic swelling of ependymal tanycytes in AD-affected hippocampus

Comparison between AD-unaffected and AD-affected hippocampus demonstrates that structural abnormalities in the latter are consistently associated with swelling tanycyte processes and associated swell-bodies. Structural differences between tanycyte processes in the alveus are particularly apparent in anti-tau labeled and Luxol H&E-stained preparations (Fig 10A-F). In non-AD-affected tissue tanycytes and swell-bodies with associated tanysomes often project in parallel fashion. Although tanysomes form receptacles, they are only weakly stained for tau-protein (Fig 10B, C). In contrast, AD-affected tanycytes appear spongiform, due to the formation of numerous circular structures and bulging waste receptacles that emanate from tanysomes (Fig 10D-F, M, N). The heavily myelinated and swelling nature of these tanycyte-derived ring structures is evident in Luxol blue H&E-stained brain sections (Fig 10D, inset, M). The comparatively strong Tau-immunolabelling in AD-affected brain tissue is predominantly associated with tanysomes, swelling toroids and associated receptacles (Fig 10D-E, O, P). Similar hypertrophic abnormalities can be observed in swell-bodies (Fig 10I-L), receptacles that project into neurons (Fig 10G, H) and tanycyte processes (Fig 10M, N, O, R). We repeatedly observed tanycyte swelling associated with bulging receptacles in the vicinity of brown deposits along tanycytes, the appearance of which is consistent with cellular waste (Fig 10J).

**Fig 10.**
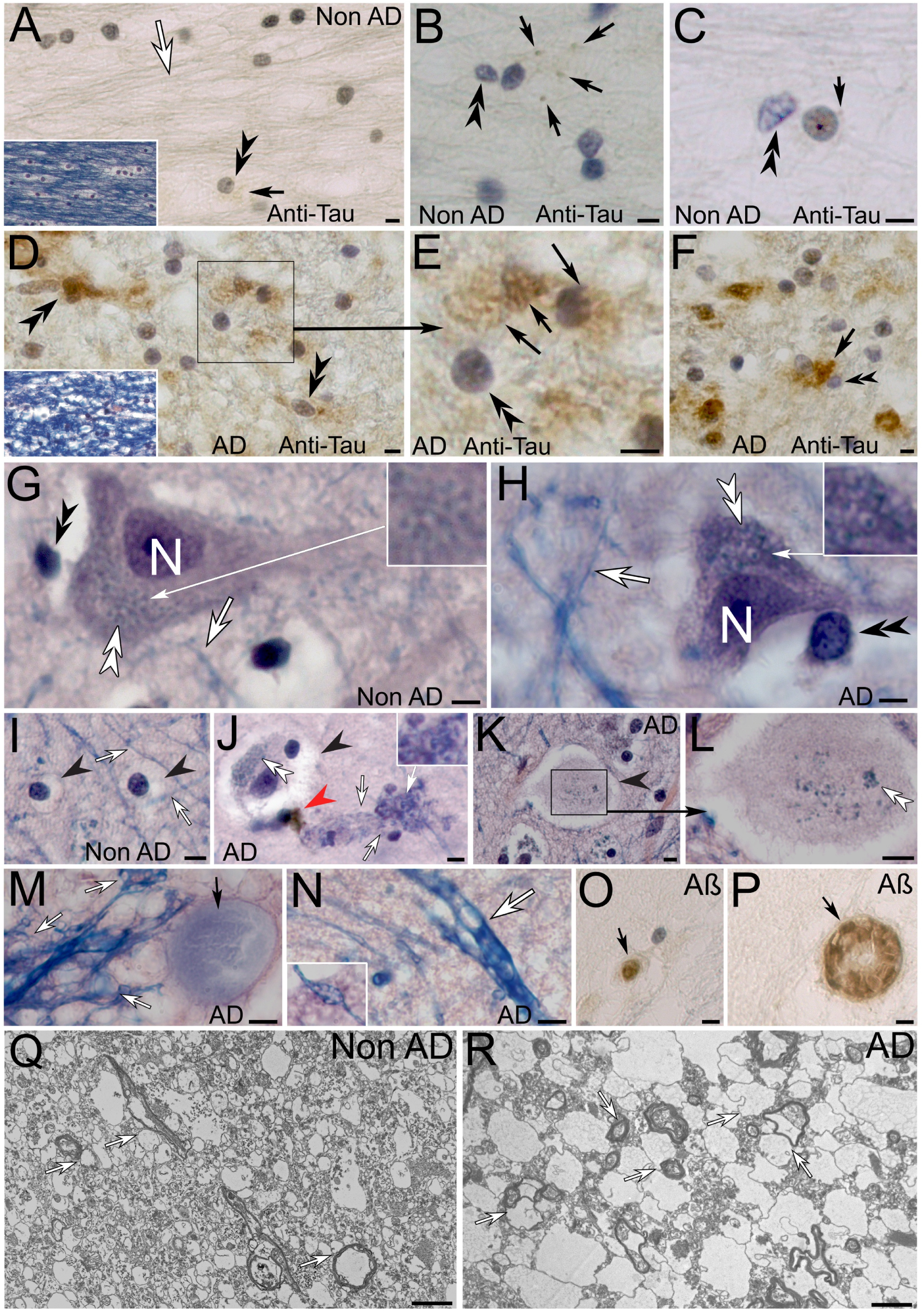
Hypertrophic tanycyte swelling in human AD-affected brain tissue. **(A-C)** Tanycyte processes in the alveus (*white arrows*) in non-AD-affected brain appear linear and project parallel to each other in both anti-tau labeled brain tissue (*A-C*) and Luxol H&E-stained preparations (*inset in A*). Tanysome (*black double arrowheads*) derived waste receptacles are sparse and appear either unlabeled or weakly labeled for tau protein (*black arrows*). **(D-F)** The alveus in AD-affected brain tissue appears spongiform with bulging tanycyte processes in both anti-tau (*D-F*) and Luxol H&E-stained preparations (*inset in D*). Numerous swelling anti-tau labeled waste receptacles (black arrows) emanate from associated tanysomes (*back double arrowheads*). **(G, H)** Similar hypertrophic swelling can be observed in tanycyte processes (*white arrows in G compared to H*) and intraneuronal tanycyte receptacles (*white double arrowheads in G compared to H, higher zoom of receptacles in respective insets*). **(I-N)** Swell bodies and associated tanycyte profiles in non-AD-affected tissue appear comparatively small and intact (*I*) compared to those in AD-affected brain tissue (*J-N*). In the latter swell-bodies, tanysome-derived waste receptacles and associated tanycytes show abnormal hypertrophy (*J-L*). Please note the accumulation of brown deposit within the hypertrophic tanycyte process, indicative of a blockage by cellular waste (*red arrowhead in J*). Tanysomes and associated waste receptacles are difficult to recognize in swell-bodies that show advanced swelling stages (*K, L higher zoom that visualizes remaining waste receptacles*). Hypertrophic tanycyte processes form numerous swelling circular structures in comparison to those in non-AD-affected tissue (*arrows in I compared to M, N, inset in N*). **(O, P)** Differences in swelling patterns are also observed in Aβ immunolabeled tanycyte-derived toroids (*arrows in O and P*). **(Q, R)** Comparison between non-AD-(*Q*) and AD-affected tissue (*R*) at the ultrastructural level demonstrates abnormal spongiform swelling that originates from bulging myelin-derived tanycyte protrusions, (*white arrows in Q compared to R*). Scale bars: A-P: 5 µm, Q, R: 2 µm.

### mRNA gene expression is indicative of the existence of different ependymal cell types in the human hippocampus that may form synergistic interactions

To test whether the myelin-forming tanycytes also express AQP4 we have conducted RNAscope experiments for AQP4, MBP and GFAP. Interestingly our findings suggest that at least two distinct tanycyte populations may contribute to the formation of the myelin/AQP4 double labeled tanycyte processes shown here (Fig 11). In the areas tested we observed that the outermost ependymal cells in the alveus expressed mRNA for both AQP4 and GFAP but lacked a signal for MBP. These observations are consistent with the Luxol H&E stain that shows lack of myelin in the alveus (Fig 11 inset in A). However, gene expression in the stratum oriens shows partial co-localization of all tested mRNAs. These findings are consistent with the hypothesis of diverse ependymal cells that may have close synergistic interactions. Our observations of anatomically different myelin-forming ependymal tanycytes in the ventricular lining that are closely associated with each other further support this postulation (Fig 11E-G). It is furthermore supported by Luxol H&E stained myelinated tanycyte processes that show varicose swellings. Close examination of these processes shows slender, blue stained processes that appear interconnected with more transparent, slightly pink stained processes clearly visible in Fig 11 panels H-J. In this profile, the myelinated part of the tanycyte forms translucent receptacles that appear unstained in this preparation and gives the tanycyte a ‘fractioned’ appearance (white arrow in H), whereas a slightly pink stained process is continuous within the tanycyte process (Fig 11I, J). These findings suggest that we may have to look at myelinated cell processes as synergistic units of diverse cell types that closely interact with each other (indicated by the red rings in Fig 11 K-M) rather than individual cell processes. Consistent with this postulation are observed close interactions between neighboring cell profiles at the ultrastructural level (Fig 11 K green arrowhead). Such synergistic interactions would explain (*i)* why myelinated tanycytes show double labelling for both AQP4 and myelin whereas mRNA expression distinguishes these two cell populations, and (*ii*) why myelinated tanycytes form varicose swellings (Fig 11G) that are likely formed by the AQP4-expressing cells. As discussed below, we have described similar synergistic interactions between AQP4 and myelin forming cells in a recently discovered waste removal system in spider brain.

**Fig 11.**
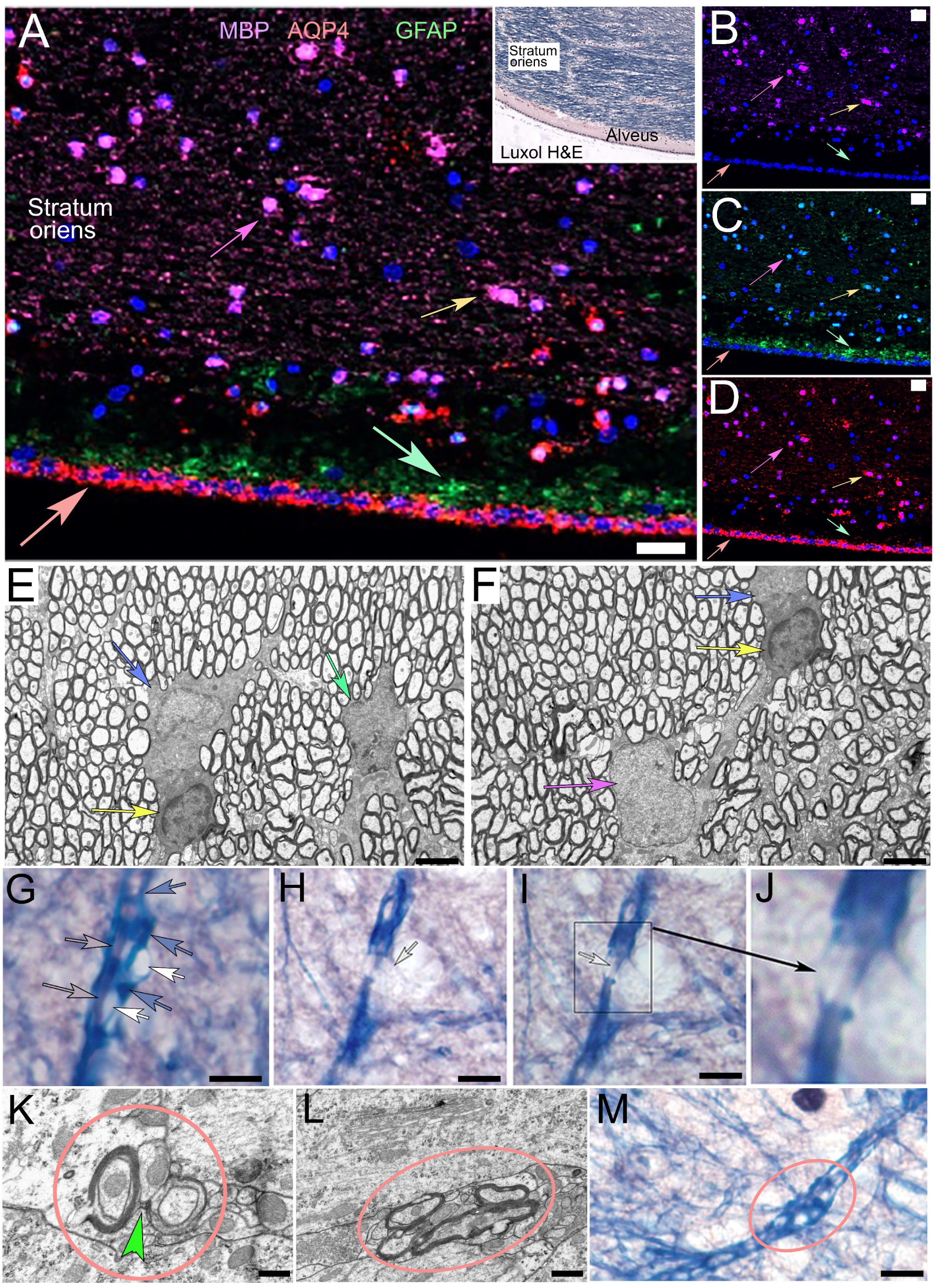
Diverse types of ependymal cells in human and rat hippocampus. **(A-D)** RNAscope labeling for myelin basic protein (*magenta arrows*) Glial fibrillary acid protein (*GFAP, green arrows*) and Aquaporin 4 (*AQP4 red arrows*) demonstrates mRNA expression for GFAP and AQP4 in the alveus, where no signal for MBP was detected consistent with Luxol H&E-stained human hippocampus (inset in A). However, all three mRNA signals partially overlap in the stratum oriens (*yellow arrow*) that contains numerous swell-bodies. (E-F) Electron-micrographs of myelin-forming ependymal tanycytes that appear anatomically different (indicated by blue, green and pink arrows) based on their nuclei and the density of their cytoplasm. Please note the close proximity of these cells with each other. (G-J) High magnification of myelinated tanycyte processes indicated by the Luxol H&E-stain. (G) Close examination shows that three different structures can be identified within this myelinated process: (i) a light, pink-stained process (*pink arrows*), (*ii*) varicose-translucent swellings (*white arrows*), and (iii) the myelinated process that has a reticular appearance (*blue arrows*). The tanycyte process in *H-J* appears ‘fractioned’ at first, however close examination shows the formation of translucent receptacles that emanate from the tanycyte (*white arrow in H*). Alteration of the focal plane reveals that the contained light pink cell process within the main tanycyte is continuous with the lower part of the tanycyte process (*white arrow in I, higher zoom in J*). These observations suggest that varicose tanycyte processes may not consist of only one cell process but may consist of diverse cell processes that form synergistic units. **(K, L)** Electron micrographs of closely associated cell profiles including myelinated cell processes. Please note the physical interconnection between two cell profiles in K (*green arrowhead*). **(M)** Varicose Luxol H&E stained tanycyte process that contains both blue and lightly pink stained cell processes. **Scale bars:** A-D 20 µm; E, F: 2 µm; G-I: 1 µm; K, L: 500 nm; M: 2 µm.

## Summary

Based on the findings presented here, we propose that the synergistic interaction between myelin-forming and AQP4-expressing ependymal tanycytes give rise to an extensive, interconnected network of myelinated cell processes that internalize and expel waste from the brain (Fig 12). Based on our findings we postulate that swell-bodies that form along the tanycyte processes represent dynamically renewable, discrete waste-internalizing units that contain the mRNA required for the formation of functional waste-internalizing receptacles. This mRNA is likely anchored within swell-bodies by mRNA-binding proteins of the quaking family. We predict that receptacle differentiation is triggered in response to (currently unknown) biochemical signals, and that AQP4-, GFAP-, Pres1, APP, tau, Sox 10, quaking, caspase 2 and 3 and Aβ are proteins that play important functional and structural roles in the differentiation of waste receptacles. We propose that receptacles project into both cell somata and extracellular spaces. Based on our preliminary functional experiments, we predict that waste uptake is mediated by AQP4, likely through the formation of a convective flow toward the receptacles. The observed Aβ-immunoreactive nature of waste-receptacles in both healthy and AD-affected human hippocampus led us to suggest that the functional role of Aβ may be the structural stabilization of the membranous receptacles to prevent their collapse during waste-uptake (see discussion). We propose that caspases 2 and 3 may play important roles in waste catabolism and postulate that hypertrophic swelling of this tanycyte-derived waste canal system leads to obstruction and depletion of neuronal somata that contain tanycyte-derived receptacles culminating in gliaptosis (macroglia-induced cell death).

**Fig 12.**
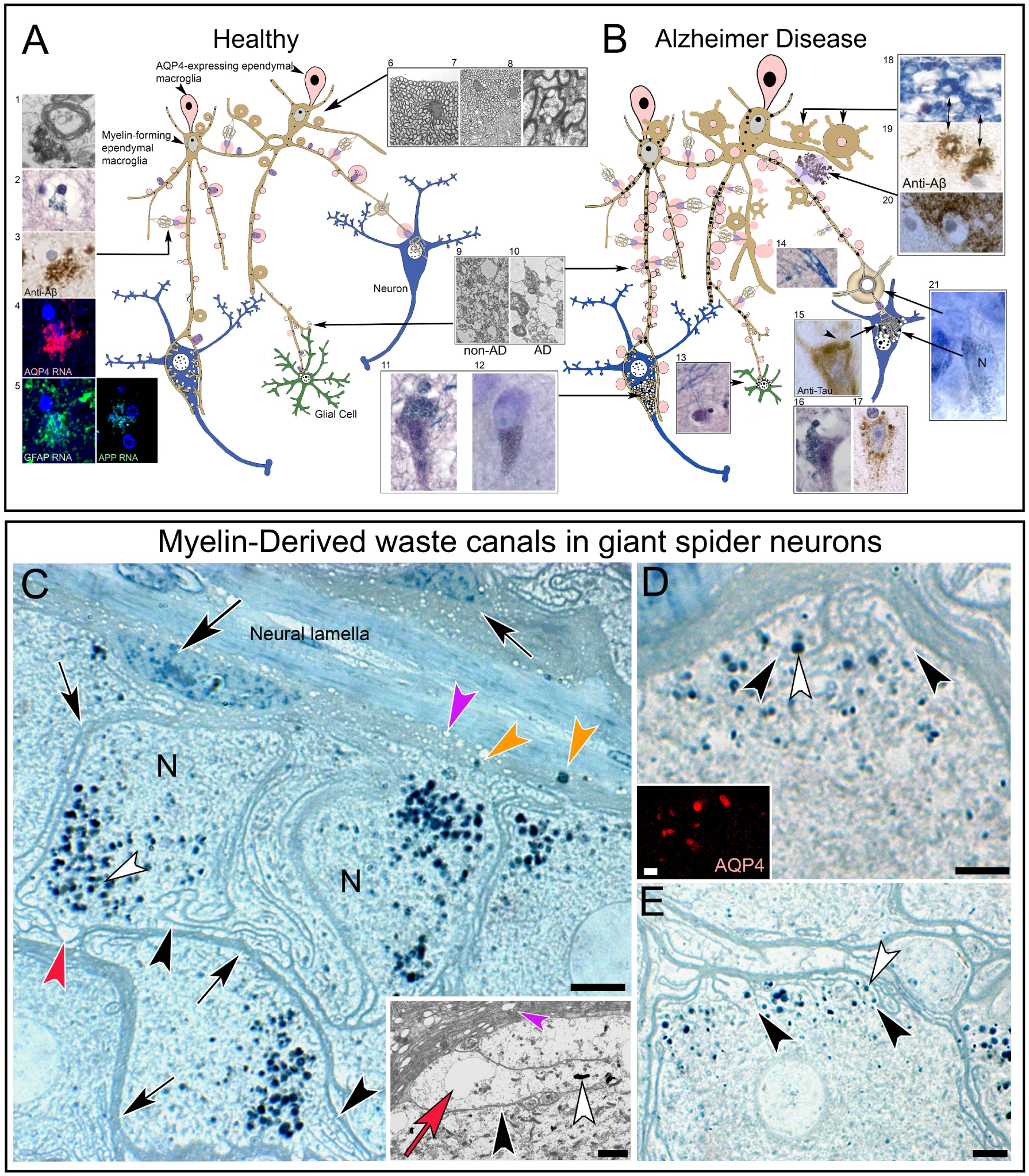
Depiction of proposed tanycyte-derived waste canal system in the human and spider brain. **(A, B)** Healthy and AD-affected brain tissue neurons (*blue*), and glial cells (*green*) are contacted by synergistically interacting aquaporin-4-expressing (light pink) and myelin-expressing ependymal tanycytes (*beige*). We postulate that interactions between the two cell types result in the formation of waste-internalizing receptacles that emanate either directly from tanycyte protrusions (inset *1) or differentiate within swell-bodies (inset 2-5*). Our findings show that receptacles that emanate from swell-bodies project into neuronal somata where they form structures currently known as ‘lipofuscin’ (*inset 11, 12*) We postulate that waste uptake takes place in an AQP4-dependent manner and that GFAP, myelin and Aβ contribute to the structural formation and stabilization of receptacles. **(B)** In AD-affected hippocampus, hypertrophic tanycyte abnormalities that may be caused by physical waste obstruction may cause swelling of the tanycytes affecting the brain parenchyma (*9, 10*). This swelling that also affects intracellular waste receptacles causes dense obstruction of affected neurons and glial cells (*11, 12, 13*), tanycyte processes (*14*), and intraneuronal tanysome-derived receptacles (*15, 16, 17*). Swelling of tanycytes in the alveus and increased intracellular turgor likely leads to excessive ‘sprouting’ of tanycyte-derived waste receptacles that appear Aβ immunoreactive as have the appearance of Aβ plaques (*18, 19, 20, 21*). **(C-E)** Histological- and ultrastructural depictions of neurons in leg ganglia of *Cupiennius salei*. Please note the large amount of cellular waste produced in metabolically active neurons and the strict containment of this waste within neurons (*white arrowheads*) or glial cells (*orange arrowheads*). Myelin-forming macroglia (*white arrows*) that contain electron-lucent inclusions (*pink arrowheads, inset showing the ultrastructural level*) line the outer circumference of each ganglion and form long myelinated projections deep into the neuronal tissue (*black arrows in C*). Each neuron contains myelin-derived canals that branch from the glial cell (*black arrowheads*) and accumulate electron-dense lipofuscin (*white arrowheads*). (*Red arrowhead in C*): swelling electron-lucent cell profile near waste particles. *(Inset in C)* Ultrastructural architecture of a myelin-derived, waste containing (*white arrowhead*) canal formed by detached myelin-membranes (*black arrowhead*). Please note the swelling translucent membrane-bound profile in the lumen of the canal (*red arrow*). (*Inset in D*): AQP4-immunoreactive nature of the electron-lucent inclusions in the myelinated glial cells (*pink arrowheads in C, inset*). Please note the unstained nature of the myelinated glial cells. **Scale bars:** C: 8µm, Inset: 2 µm; D: 1 µm, Inset 5 µm; E: 5 µm. Schematics are not drawn to scale.

## Discussion

The findings presented here are consistent with our previous postulation that myelinated tanycytes reside in the ventricular lining and form varicose, slender processes that project into the hippocampal formation and form receptacles that project into neurons and extracellular spaces (see also [10]). Based on the findings presented here, we argue that these cells are different from cells that are currently described as oligodendrocytes for the following reasons: (1) Mature myelinating oligodendrocytes are believed to be post-mitotic in healthy brain [29]. Thus our observations that show that the ependymal cells are often multinucleated although they are clearly forming myelinated processes make it unlikely that these are dividing, myelinating oligodendrocytes (see also [10]). (2) Both cell anatomy and location of described oligodendrocytes are different from the myelin-forming macroglia shown here. Currently, oligodendrocytes are described throughout the brain parenchyma [30], whereas the somata of the myelin-forming macroglia shown here appear strictly contained in the ependymal lining with only their myelinated cell processes projecting into the hippocampal formation. (3) The myelin-forming macroglia form a dense, interconnected reticulum similar to our observations of myelin-forming waste internalizing macroglia in spider brain shown here. These observations are inconsistent with our current understanding of oligodendrocytes in healthy brain tissue. We therefore postulate that the ependymal myelin-forming and AQP4-expressing cells with their long processes that project deep into the brain parenchyma are likely previously unrecognized tanycytes. We concur with Pasquettaz et al 2021 who also described ‘peculiar protrusions’ along tanycyte processes, that a far greater number of tanycytes may exist than currently recognized [13]. Although we have only investigated mammalian hippocampus in this study, it would not be surprising if other brain areas show similar myelin/AQP4 expressing macroglia that project from the ependymal lining into the brain parenchyma and form waste-receptacles consistent with lipofuscin [31–34]. This study demonstrates the monumental importance to re-investigate the histological and functional role(s) of myelin in the brain using modern technology that far exceeds the tools available to early pioneers that established our current understanding of myelination in the 19^th^ and 20^th^ centuries [35].

### How can AD-related histopathologies be explained in context of the proposed waste-internalizing glial canal-system demonstrated here?

#### The proposed synergistic interaction of myelin-forming and AQP4-expressing ependymal tanycytes

First, we would like to emphasize that we do not claim that our below-mentioned postulations are ‘proven fact.’ We strive to propose a logical and testable model based on our supportive findings and other published observations cited below.

Based on our studies in spider, rodent, and human brain tissue, we propose the existence of a waste-internalizing glial canal system in the brain that provides a logical explanation for: *(i)* controlled waste removal from the brain; *(ii)* the proposed connection between ependymal macroglia, astrocytes, and oligodendrocytes; *(iii)* the proposed underlying causes for reactive astrocytes and oligodendrocytes; *(iv)* common histopathologies described in AD-affected brain; and *(v)* proposed underlying causes for these pathologies.

##### (i) Controlled waste removal from the brain

Waste removal in the human body is essential for survival and is governed by specialized canal systems. These systems are highly conserved throughout the phylogenetic tree as they fulfil the life-sustaining function of detoxification by processing, sequestering, and expelling endogenous metabolic end-products and exogenous xenobiotics (e.g., ingested toxins) [36]. In mammals, such waste canal systems include the lymphatic, urinary, cardiovascular, pulmonary, and hepatic portal systems. What these systems have in common is that they are: (a) closed-loop transport systems that strictly sequester waste without spillage and expel this waste from the body, (b) biochemically governed and structurally stabilized uptake mechanisms that effectively internalize waste products without collapsing, (c) interconnected, and (d) prone to obstruction or rupture of the canal systems, which may be fatal if untreated [37].

Although neurons are among the most metabolically active cells in the human body that necessarily create large amounts of metabolic end products [22, 38, 39], we currently have no clear understanding how this waste is contained, catabolized, and expelled from the brain. One would argue that it is ***especially important*** to strictly contain both lipophilic and hydrophilic waste in the brain to avoid unspecific interactions of waste with both intracellular and extracellular domains of ion channels, transmembrane transporters, transmitter receptors, transcription factors, and other important proteins, without which signal transduction in the nervous system would rapidly fail. Such unspecific interactions are demonstrated by highly lethal inhaled (sarin gas; [40]), injected (α-bungarotoxin;[41]) and ingested (tetrodotoxin; [42]) toxins and venoms. It is this aspect of the glymphatic-hypothesis, the proposed free release of cellular waste into the brain parenchyma that we respectfully put into question [1, 5–7]. Giant neurons in the spider brain in which cellular waste is clearly visible demonstrate the large amount of waste that is produced in neurons. Although this waste is less conspicuous in the much smaller mammalian neurons, nevertheless the increased numbers of neurons in the vertebrate brain together with narrow interstitial spaces would likely also pose the risk of both lipophilic and hydrophilic waste aggregates that would be challenging to clear from extracellular spaces. Finally, waste-clearing canal systems in the body are highly conserved throughout evolution due to their importance for survival, examples include the urinary [43] or intestinal tracts [44]. We therefore postulate that the myelin-derived waste canal system we have discovered in giant arachnid neurons [10], although structurally altered is likely highly conserved in the mammalian brain; what is particularly compelling is the observed and likely synergistic interaction of myelin and AQP4. We propose that it is this interaction that likely provides the renewable structural foundation of waste containment (myelin), and a driving force that draws cellular waste out of neurons into the canal system (AQP4) [10, 22]. We recognize the limitations of our preliminary functional uptake studies in mouse brain. However, we have previously conducted similar, successful experiments in spider brain to locate the AQP4-expressing cell profiles we detected in the lumina of myelin derived waste canals. The myelin-forming macroglia did not show AQP4 immunolabeling, suggesting the synergistic interaction between two different cell types. Unable to find the AQP4 immunoreactive cells in our brain sections, we conducted a similar uptake experiment using methylene blue vital dye. After 30 min, we discovered the ‘blue cells’ in the dorsally located tubular system that we routinely removed from our brain preparations. This is why we selected a similar timeframe in our mouse experiment. Interestingly, Sauvé et al. (2026,) utilized a similar timeframe for their functional studies that enabled them to successfully show internalization of tau protein by tanycytes [12]. Immunolabeling confirmed the AQP4-immunoreactive nature of these cells and their long, slender processes that project into the spider brain [10]. We therefore tested this method in mouse brain. Based on the immunohistochemical, histological, and ultrastructural experiments, we predicted that the following structures would internalize fluorochrome and therefore fluoresce: (i) the ependymal layer and alveus, (ii) tanycyte processes that project into the brain, (iii) swell-bodies and associated receptacles, and (iv) drainage canals that channel waste from the brain. We predicted that neurons would not fluoresce due to lack of penetration of these fluorochromes into living cells. All of these predictions were observed in our experiments, some of which are shown here. This experiment was designed to test the general validity of our hypothesis and will serve as guidance for more targeted functional studies moving forward. We decided to include these experiments in order to show other scientists who may be interested in conducting functional experiments which structures can be expected to internalize waste and should therefore be included in their preparations.

##### (ii) The proposed connection between ependymal macroglia, astrocytes, and oligodendrocytes

As shown here, the myelin-forming ependymal macroglia are distinct from oligodendrocytes with respect to both cell anatomy and spatial distribution in the mammalian brain. However, we postulate that these two structures are formed by the synergistic interactions of (at least) two types of ependymal macroglia that give rise to both receptacles and swell-bodies along their interconnected cell processes. These cell types are: (i) the AQP4 and GFAP-expressing macroglia shown here, and (ii) the myelin-forming tanycytes described here. Based on our observations in arachnid and mammalian brain, we postulate that fine cell processes of the AQP4 expressing macroglia are tightly interconnected with the cell processes of myelinated macroglia. We postulate that swell-bodies are formed by the interactions between both cell types whereby the swelling and electron-lucent parts of the swell bodies are formed by the AQP4-expressing glial cells and the myelin-derived waste receptacles that emanate from tanysomes are contributed by the myelinated macroglia. This would explain why early swell-bodies that have not yet formed receptacles show a stronger signal for myelin and a less prominent signal for AQP4, since this water channel is likely only expressed at higher levels once the functional receptacles form. In contrast, the myelin that we propose is utilized for the structural formation of functional receptacles and may be supplied through the tanysome connection with the tanycyte process is weaker (‘used up’) in swell-bodies with formed receptacles.

##### (iii) The proposed underlying causes for reactive astrocytes and oligodendrocytes

We postulate that swell-bodies whose tanysomes do not form receptacles represent ‘dormant,’ stage I [28] ‘waste containers’ similar to waste containers we distribute throughout cities and buildings to collect, contain and remove waste. We postulate that activity-, stress, and waste-related signals (e.g., transmitter receptors, cytokines, hypoxia, ion gradients) activate dormant swell-bodies and that this activation leads to AQP4- and likely ion-channel mediated water- and ion-influx, resulting in changing concentration gradients and altered gene expression. To this end, the observed synergistic interactions between AQP4 and the swelling-sensitive transient receptor potential isoform 4 (TRPV4), need to be investigated [45]. Such mechanisms are well established in the development and differentiation of cell structure [46] and would logically explain: (i) the electron-lucent nature of swell-body lumina and their different swelling patterns [28], (ii) the physical connection of tanysomes with tanycyte processes to ensure the continued supply of essential building blocks for the continued formation of fully differentiated and functional receptacles, (iii) the presence of multiple nucleus-like tanysomes and toroids in individual swell bodies that may be indicative of increased brain activity that requires the dynamic differentiation of additional waste receptacles to ensure effective waste uptake, and (*iv*) the presence of mRNA-binding proteins of the quaking family in receptacle forming toroids, as they require the building blocks and blueprints for dynamic and rapid ‘on-site’ receptacle differentiation where metabolic activity or injury-related cellular waste is formed. Furthermore, this proposed model would also explain why the tanysomes express the Sox10 gene that is absent in emerging toroids as shown here. We postulate that Sox10 likely prevents the differentiation of active tanysomes into waste receptacles and enables them to give rise to additional differentiating toroids that lack Sox10 expression and therefore differentiate functional waste receptacles. We postulate that Sox10 is downregulated in aging swell-bodies that lose their ability to form additional toroids and that aging toroids form waste-receptacles instead. Mechanisms that trigger the downregulation of Sox10 could be related to the gradual depletion of vital proteins that are essential for the formation of waste-receptacles. Such concentration gradient related mechanisms are well described in the development of structure. Based on observations that are part of our ongoing research, we postulate that aging swell-bodies detach from the tanycyte processes, similar to ripe apples that detach from tree branches and are channeled out of the brain toward the ventricles. The underlying mechanisms for this proposed model are subject to our ongoing research.

##### (iv) Common histopathologies described in AD-affected brain

Among the most prevalent histopathologies reported in the human AD-affected brain are Aβ plaques, fibrillary tau tangles, spongiform abnormalities, senile plaques, and neuronal obstruction with lipofuscin [47–50]. As demonstrated in this study, these pathologies can be logically explained if we put this proposed waste-canal system in place and consider that it might be physical obstruction of this canal system that leads to hypertrophic swelling of its associated structures.

###### Spongiform abnormalities

Such swellings can be explained by the swelling of the AQP4-expressing macroglia in this synergistic system. As shown here, forming receptacles appear almost transparent in Luxol H&E-stained preparations, which is why they are easily missed when not actively searching for these receptacles. These translucent structures are why swelling of this system may be mistaken for ‘empty’ spaces, rather than hypertrophic receptacles.

###### Neuronal obstruction by lipofuscin

We have demonstrated at both the light- and electron-microscopic levels that myelinated glial-processes form bead-like varicosities that project into neuronal somata where they differentiate into waste-internalizing receptacles known as lipofuscin in both non-AD-affected and AD-affected brain [10, 28]. Lipofuscin has been described as insoluble and of uncertain origin [51]. The insoluble nature of these receptacles would be explained by the hypothesis that lipofuscins are dynamic myelin-derived glial projections that consistently internalize and dispose of cellular waste in healthy neurons. This postulation is consistent with our observations of myelin-derived processes that project into neuronal somata at both light and electron-microscopic levels [10]. Such projections stain prominently for Luxol blue in paraffin-embedded sections of AD-affected human hippocampus [10]. Blockage-related swelling of this glial-canal system would therefore explain why intraneuronal tanycyte processes obstruct neuronal somata in AD-affected brain.

Interestingly, strongly Luxol blue stained neuronal somata have also been described in the brains of cats, dogs, and mice with neuronal ceroid lipofuscinosis (NCL) [10, 52, 53]. Moreover, Luxol blue stain is also observed in neurons of the human variant of NCL known as Batten disease [54, 55]. In humans, this rare early onset genetic disorder is linked to mutations in ‘Ceroid Lipofuscinosis, Neuronal genes CLN1-14’ a family of genes often linked to catabolic lysosomal enzymes [54, 56]. One of the most prevalent mutations in Batten disease affects palmitoyl-protein thioesterase 1, an enzyme that cleaves palmitate (16-carbon fatty acid chains) from proteins. Such lipid anchors are abundant in neurons due to their ability to dynamically control protein trafficking and localization of vital proteins at synaptic sites [57, 58]. It remains to be investigated whether/how this inability to cleave palmitate from damaged proteins is related to the prominent myelin-stain within neuronal somata. It cannot be excluded that the attached lipid chains prevent the internalization of damaged proteins by myelin-derived waste receptacles and instead anchor damaged proteins in the receptacle membrane leading to impairment of the waste-canal system, and rapid buildup of cellular waste in affected neurons.

###### Amyloidβ plaques

We demonstrate that receptacles that emanate from tanysomes contain Pres1 and APP mRNA. We further show that this mRNA is likely expressed by the AQP4-expressing ependymal cells that show gene-expression and immunoreactivity for both genes and their encoded proteins. Waste-internalizing systems require physical stabilization of the waste-internalizing structures to prevent their collapse. Numerous examples of such specialized structures are described in both the human body (e.g., pulmonary system stabilized by cartilaginous rings [59]) and human technology such as stabilized vacuum hoses or dryer vents. The proposed waste internalization presented here consists of membranes that form the proposed canal structures, which are clearly visible in giant spider neurons. To prevent that these membranes get sucked toward the proposed AQP4-generated convective flow, such a system requires structural stabilization. Interestingly, in both non-AD- and AD-affected human brain the proposed waste-internalizing receptacles are consistently associated with Aβ. Research into this protein reveals that it is a hydrophobic protein that would therefore interact with membranes and forms structurally very stable beta-pleated sheets [60, 61]. These characteristics of Aβ42 make this protein a suitable candidate to stabilize waste internalizing structures. Here, we demonstrate that Aβ-plaques consist of receptacles that show abnormal proliferation and hypertrophy in AD-affected brain. These findings are inconsistent with the current amyloid hypothesis that postulates the random accumulation of misfolded Aβ42 that has - despite massive international research efforts - failed to illuminate the underlying causes for this disease. Moreover, it would also explain, why the removal of Aβ using monoclonal antibodies has repeatedly failed in drug trials, since the lack of this stabilizing protein would result in the collapse of remaining healthy waste-internalizing tanycyte processes and likely interfere with water flow in the brain [62–64]. Lastly, our postulation suggests a vitally important role of Aβ in waste removal from the brain. This would explain why familial Alzheimer cases are strongly associated with mutations in the presenilin1, presenilin2 and APP genes [65]. All three genes play vitally important roles in the synthesis of Aβ [66]. Based on our postulation, the waste-internalizing structures in these patients are likely not sufficiently stabilized.

Due to our findings that the Aβ42 labeled areas show clear structure that we have identified as waste-internalizing receptacles through correlative light and electron microscopy, we support the growing notion that the amyloid hypothesis should be rejected [67–69] and alternative, more consistent explanations should be explored, one of which is presented here.

Other abnormalities observed in AD-related proteins are likely important for a multitude of different structural and biochemical processes required in such a system. A human metaphor comparison is the emptying of a septic tank that requires: (a) an opening in the septic tank (possibly, alpha synuclein; [70]), (b) a hose that can be lowered into the septic tank to internalize the sewage (myelinated tanycyte processes that project into neurons); (c) a pump that applies appropriate suction to remove the sewage from the septic tank into the hose (AQP4-mediated convective flow); (d) a septic truck tank receptacle that contains and transports the sewage for proper disposal (swell bodies with accumulated waste); a sewage processing plant that utilizes chemicals to process the sewage (caspase 2, 3 and other aforementioned catabolic enzymes); and the treated waste is disposed of into local waterways (ventricular release). In this example, only a few required components are listed. One can argue that the building and maintenance of the required equipment and processing plants are also important in this process. Such mechanisms are commonly utilized in human technology and throughout the human body and would explain the abundance of proteins that are implicated with AD [71].

##### (v) Proposed underlying causes for these pathologies

What all these systems have in common is that they sometimes fail for numerous reasons, from blockage to equipment failure, and it is thus not unreasonable to postulate that AD may be caused by the blockage of this proposed synergistic waste-canal system. The prevalent comorbidity of AD with fungal, viral, and bacterial infections provides a compelling explanation for the blockage of a narrow glial canal system [72–74]. Fungal and bacterial cell walls or structural viral proteins may not be catabolized as efficiently as endogenous metabolic waste and may result in obstruction. Yet, critics of our hypothesis reject that such a commonly utilized waste-internalizing canal system may exist in the brain to ensure safe and controlled waste removal from the brain, one of the most metabolically active organs in the human body. After decades of testing the amyloid hypothesis without success, we encourage the scientific community to not outright reject the glial-canal hypothesis that was guided by the clear visibility of this system in giant spider neurons. We encourage scientific discussion and for other groups to consider the existence of this system in their research and either support or refute some or all of this hypothesis.

## Conclusions

We acknowledge that the glial-canal-hypothesis presented here contradicts current understanding of structure in neuroscience. We have thus provided numerous example picture panels to support our statements. We have countless additional supportive experiments, images, and data that are too extensive to include in one manuscript. We are happy to provide to the neuroscience community access to our preparations, technology, data, and images and strongly encourage the independent re-investigation of cellular structure in the human and mammalian nervous system by other investigators with their own samples. We suggest that particular focus should be given to the identity of myelinated cell profiles using either longitudinal sections through neurons as presented here, or clear ultrastructural serial section analysis to unambiguously demonstrate the origin and identity of myelinated cell processes in the brain.

## Acknowledgements

Research reported in this publication was supported by an Institutional Development Award (IDeA) from the National Institute of General Medical Sciences of the National Institutes of Health under grant number P20GM103449. Its contents are solely the responsibility of the authors and do not necessarily represent the official views of NIGMS or NIH. We thank Dr. John DeWitt for support and access to human decedent tissue. We thank Drs. Alain Brizard, Christopher Francklyn, Mark Nelson, Mark Lubkowitz, Douglas Taatjes, and Heather Driscoll for support and helpful feedback on this work. We thank Ann and Dr. David MacLaughlin for their donation to fund the Zeiss confocal microscope. The electron microscopic investigations were performed at the Microscopy Imaging Center at the University of Vermont (RRID:SCR_018821). We thank Dr. Gerald Herrera, Nicole Bouffard, Kyra Lee, and Brad Vietje for technical support.

## Abbreviations

AQP4: Aquaporin-4 aqua channel
AD: Alzheimer Disease
Aß: Amyloid beta
APP: Amyloid Precursor Protein
GFAP: Glial Fibrillary Acid Protein
IR: Immunoreactive
NCL: Neuronal Ceroid Lipofuscinosis
Pres1: Presenilin-1

## Notes

### Competing Interest Statement

Dr. Adam Weaver and Abigail Roman declare no competing financial interests.
Saint Michael's College, together with Dr. Ruth Fabian-Fine, have filed a patent as a direct result of this research.

### Summary of Updates

We needed to revise to meet additional Version of Record and MDAR requirements for the Journal eLife.

